# Integrated Multiomics Links Metabolic and Inflammatory Remodeling to Arterial Stiffness After the 4,486-km Trans Europe Footrace

**DOI:** 10.64898/2026.03.12.711477

**Authors:** Christopher M Clements, CeAnn C Udovich, Katelyn R Ludwig, Francesca Cendali, Monika Dzieciatkowska, Sotirios P Fortis, Uwe H Schütz, Arno Schmidt-Trucksäss, Christopher Klenk, Angelo D’Alessandro, Douglas R Seals, Zachary S Clayton, Travis Nemkov

**Author notes:** **<u>Corresponding authors:</u>** Travis Nemkov, PhD, Department of Biochemistry and Molecular Genetics, University of Colorado Anschutz Medical Campus, 12801 East 17th Ave, Mail Stop 8101, Aurora, CO 80045, Zachary Clayton, PhD, Department of Medicine-Geriatrics, University of Colorado Anschutz Medical Campus, 12631 East 17th Ave, Mail Stop B179, Ste 8111, Aurora, CO 80045. These authors contributed equally and share the first authorship (Christopher M Clements, CeAnn C Udovich, Katelyn R Ludwig).

## Abstract

**Rationale:** Regular aerobic exercise protects against vascular aging and reshapes the circulating molecular milieu, but the relation between vascular function, circulating molecules, and exercise dose at extreme volumes remains poorly defined. The vascular and molecular consequences of chronic, multi-stage ultra-endurance running are particularly unclear.

**Objective:** To define circulating molecular signatures associated with vascular dysfunction following the 64-stage, 4,486-km Trans Europe Foot Race (TEFR).

**Methods and Results:** Integrated multiomics analysis (proteomics, lipidomics, metabolomics) of plasma from 27 finishers revealed a coordinated systemic shift driving an oxidative phenotype. Specifically, we identified altered arginine metabolism and a universal upregulation of lipotoxic ceramides consistent with incomplete fatty acid oxidation. In conjunction, we identified upregulation of innate immune system pathways including the acute phase response and the complement system.

Central pulse wave velocity (cPWV) increased significantly after the race, consistent with arterial stiffening. To test whether the post-race circulating milieu could directly influence vascular mechanics, naïve murine aortic rings were incubated with participant plasma. Post-race plasma acutely increased aortic elastic modulus, and this effect was attenuated by the superoxide dismutase mimetic TEMPOL, supporting a ROS-dependent component. In human aortic endothelial cells (HAECs), post-race plasma increased reactive oxygen species generation without detectable changes in eNOS phosphorylation, total eNOS abundance, or stimulated nitric oxide production. Endothelial ROS responses were associated with components of the terminal complement pathway.

**Conclusions:** Extreme multi-stage ultra-endurance exercise induces a distinct systemic milieu associated with arterial stiffening through ROS-sensitive mechanisms. This response is characterized by remodeling of arginine-related metabolism, ceramide accumulation, innate immune activation, and oxidative stress, without evidence of reduced measured eNOS abundance or stimulated NO production. These findings identify candidate molecular pathways linking prolonged metabolic stress to vascular dysfunction.

## Introduction

Vascular dysfunction – an independent risk factor for cardiovascular disease – is characterized by progressive large artery stiffening.^1^ This stiffening is driven in part by excess reactive oxygen species (ROS), extracellular matrix (ECM) remodeling, and increased vascular smooth muscle tone resulting from reduced nitric oxide (NO) bioavailability.^1^ Regular aerobic exercise is widely recognized as a cornerstone of cardiovascular health, serving as a primary non-pharmacological intervention for the prevention and treatment of vascular dysfunction associated with aging^2^. Habitual moderate-to-vigorous physical activity is generally associated with reduced arterial stiffness, improved endothelial function, and decreased cardiovascular-related and overall mortality risk^3,4^. Clinical studies have demonstrated that habitually endurance exercise-trained older adults tend to display endothelial function^5,6^ and arterial stiffness^7,8,9,10^ comparable to young sedentary adults, and that initiating aerobic training in previously sedentary older adults can improve vascular function.^11,12,13^

The relation between high-volume exercise doses and vascular health is not strictly linear^14^. Guideline-based physical activity generally maintains redox homeostasis by suppressing excess ROS-induced oxidative stress via upregulation of antioxidant defenses (consistent with hormesis), but very high, prolonged workloads can transiently amplify oxidative and inflammatory stress. Ultra-endurance running (events exceeding the marathon distance) imposes sustained mechanical, metabolic, and redox loads with limited recovery^15^. Participation in these events has expanded markedly over recent decades, particularly in midlife. During this period, the human body undergoes non-linear waves of proteostatic and organ-specific aging indicative of diminishing capacity to cope with damage and repair tissues.^16,17^ Thus, repeated mechanical, metabolic, and redox loads like those experienced in ultra distance races might accelerate vascular damage as these protective processes wane. Indeed, impaired cardiovascular health associated with ultra-distance races can persist for days or weeks after the event^18–20^, and long term high-volume exercise training has been associated with impaired cardiovascular function^21^. Thus, there is a need to better understand the impact of this stress on vascular function. This need is particularly relevant for multi-stage ultra-endurance events, where insufficient recovery between daily efforts creates a unique physiological state distinct from single-stage races.

The Trans Europe Foot Race (TEFR^22^) – a 4,486-kilometer, continent-spanning run over 64 days without notable breaks – offers a unique opportunity to characterize changes in vascular function and associated systemic alterations that occur over the span of the race. Unraveling these complex physiological shifts requires moving beyond the targeted analysis of traditional biomarkers. Therefore, the application of omics technologies such as proteomics, metabolomics, and lipidomics offers a unique ability to perform unbiased, systems-level interrogation. This multiomics approach is essential to capture the dynamic interplay between metabolic flux, lipid remodeling, and the innate immune activation that cannot be fully elucidated in isolation. In this study, we characterized samples isolated from 27 finishers before and after the 2009 TEFR to integrate comprehensive plasma multiomics with *in vivo* vascular phenotyping using central pulse wave velocity (cPWV – a non-invasive assessment of large artery stiffness) and *ex vivo* mechanistic assays using human aortic endothelial cells (HAECs) and isolated murine aortic rings. This framework enabled paired *in vivo* and *ex vivo* interrogation of how the circulating milieu relates to arterial properties and endothelial signaling during an extreme, real-world exposure.

## Methods

### Overview

Runners were followed with a mobile truck (MRI-Trailer Model Mob.MRI 02.05, SMIT Mobile Equipment B.V., Division AK Specialty Vehicles, Farnham, UK) housing a 1.5 Tesla whole-body MR imager (Magnetom Avanto™ mobile MRI 02.05, software version: Syngo™ MR B15, Siemens Ltd., Erlangen, Germany). The MRI was utilized to collect central PWV in a subset of 10 runners pre- and post-race. Blood samples were obtained pre- and post-race and immediately centrifuged to obtain the plasma component. Plasma was frozen (below -20°C) and stored in a -80°C freezer after the race^22^.

### Cell Culture Experiments

#### Reactive oxygen species production

Human aortic endothelial cells (HAECs, PromoCell; used after 4–6 passages. Donor: 80-year-old female with no cardiac diseases, non-smoker) were cultured in 96-well glass bottom plates (CellVis) under standard conditions (37.5°C, 100% relative humidity, 5% CO2) in basal media [Endothelial Cell Growth Medium-2 (EGM-2) BulletKit + 1% penicillin-streptomycin (pen-strep); Lonza] supplemented with 10% subject plasma.^23^ After a 2-hour plasma exposure, cells were stained with fluorescent probes to detect basal ROS bioactivity (5 μM CellROX Deep Red; Thermo Fisher Scientific). Cells were incubated with CellROX for 30 min, washed with HBSS, and imaged immediately. Hoechst 33342 (Thermo Fisher Scientific) was used as a nuclear stain. Images were acquired using an EVOS M7000 fluorescence microscope (Thermo Fisher Scientific). Fluorescence quantification was performed using Celleste Image Analysis Software (Thermo Fisher Scientific). These methods were conducted in all finishers (N=30).

#### Endothelial nitric oxide synthase activation

To assess the influence of circulating factors on eNOS activation for impacting NO bioavailability, HAECs treated with 10% subject plasma were washed in PBS and scraped with RIPA buffer containing protease and phosphatase inhibitors (Roche). Cell lysates were then sonicated for 20 seconds and incubated on ice for 10 minutes. Thereafter, cell lysates were clarified by centrifugation at 13,000g at 4°C for 10 minutes. The supernatant was collected and lysate concentrations were determined using a bicinchoninic acid (BCA) assay (Thermo Fisher). Relative protein amounts of phosphorylated (p)-eNOSser1177 (Cell Signaling), a primary activation site of eNOS^24^, (p)-eNOSthr495 (Cell Signaling), a primary deactivation site of eNOS, and total eNOS (Abcam), were determined by capillary electrophoresis immunoassay (Jess; ProteinSimple) Protein expression was normalized to total protein in the sample and presented as arbitrary units (AU). These methods were conducted in a subset of finishers (n=5).

#### Nitric oxide production

HAECs (PromoCell; used after 4–6 passages. Donor: 80-year-old female with no cardiac diseases, non-smoker) were cultured in 96-well glass bottom plates (CellVis) under standard conditions (37.5°C, 100% relative humidity, 5% CO2) in basal media [Endothelial Cell Growth Medium-2 (EGM-2) BulletKit + 1% penicillin-streptomycin (pen-strep); Lonza] supplemented with 10% subject plasma^23^. After a 2-hour plasma exposure, cells were stained with fluorescent probes DAR-4M AM (Sigma-Aldrich) to detect NO production. Cells were incubated with DAR-4M AM for 40 min, washed with HBSS, and incubated for an additional 30 minutes at 37°C and 5% CO_2_ before imaging. Hoechst 33342 (Thermo Fisher Scientific) was used as a nuclear stain. HAECs stained with DAR-4M AM were imaged before and 6 minutes after the addition of 100 μmol/L acetylcholine (Sigma Aldrich) to stimulate NO production. Images were acquired using an EVOS M7000 fluorescence microscope (Thermo Fisher Scientific). Fluorescence quantification was performed using Celleste Image Analysis Software (Thermo Fisher Scientific). These methods were conducted in all finishers (N=30).

### Animal Experiments

#### Ethical Approval

All mouse studies and procedures were reviewed and approved by the Institutional Animal Care and Use Committee at the University of Colorado Boulder (Protocol No. 2618). All procedures adhered to the guidelines set forth by the National Institutes of Health’s Guide for the Care and Use of Laboratory Animals.

#### Aortic Pulse Wave Velocity

Aortic pulse wave velocity (PWV) was assessed using phase-contrast cine magnetic resonance imaging (MRI), the reference standard for non-invasive quantification of arterial stiffness. All examinations were performed in head-first supine position with ECG-based prospective cardiac triggering. Total acquisition time for the central aortic PWV protocol was approximately 4–5 minutes (within a total vascular MRI protocol of ∼25 minutes).

Flow measurements were obtained at the proximal and distal thoracic aorta using a transverse 2D FLASH sequence with phase-contrast acquisition and individually adjusted velocity encoding (venc). Imaging parameters were: flip angle 20°, TE ∼2.75 ms, TR ∼11.55 ms, slice thickness 5 mm, field of view 768 mm, matrix 256 × 192, in-plane spatial resolution 1.25 mm isotropic, phase bandwidth 590 Hz/pixel, and 100 reconstructed cardiac phases per RR cycle.

Transit time between the systolic upstroke of the proximal and distal aortic flow waveforms was determined, and central PWV was calculated as the ratio of the centerline distance between measurement sites to the pulse transit time (PWV = Δx/Δt).

#### Aortic Intrinsic Mechanical Wall Stiffness (Elastic Modulus)

Aortas were promptly excised from the mice following cardiac exsanguination, rinsed with cold physiological saline solution, and cleared of any remnant perivascular connective tissue. To measure ex vivo aortic stiffness, two thoracic aorta samples (∼1 mm in length) were cut and used to determine intrinsic mechanical stiffness via wire myography as previously described^25^. Aorta samples were placed in heated (37°C) baths filled with calcium-free, phosphate-buffered saline. The samples were then mounted on two wire prongs, followed by three rounds of pre-stretching. Once pre-stretching was complete, aortic ring diameter was increased until 1 mN of force was reached and incrementally increased by 5 µm every 3min thereafter until failure. The force corresponding to each stretching interval was recorded and used to calculate stress and strain. A stress-strain curve was then generated using the following equations:

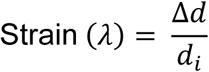

where d is diameter and di is initial diameter.

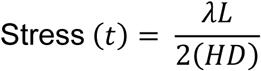

where L is one-dimensional load, H is wall thickness, and D is vessel length.

The elastic modulus of the stress-strain curve was determined as the slope of the linear regression fit to the final four points of the stress-strain curve, as previously reported^26^. To assess the stiffening role of participating in a multi-stage ultramarathon, aortas from young (3-4 month) intervention naïve donor C57BL/6 mice were prepared as described in the “Aortic Intrinsic Mechanical Wall Stiffness (Elastic Modulus)” section immediately above. Aortic segments (∼1 mm in length) from each donor animal were incubated in the following conditions in duplicate for 48 hours: 1) DMEM + 1% penicillin-streptomycin + 10% subject plasma (pre-race); and 2) DMEM + 1% penicillin-streptomycin + 10% subject plasma (post-race). The plasma samples were sex-matched to that of the donor animal. Following the incubation period, aortas were immediately mounted on two wire prongs and elastic modulus was assessed exactly as described above. “Post-race” plasma-induced changes in aortic elastic modulus were determined as the difference between post- and pre-race plasma incubation conditions. These methods were conducted in a subset of runners; those with central PWV and ROS data (n=9) and just ROS data (n=6). To determine if participating in a multi-stage ultramarathon results in arterial stiffening via excessive ROS production, aortic segments (∼1 mm in length) from each donor animal were incubated in the following conditions in duplicate for 48 hours: 1) DMEM + 1% penicillin-streptomycin + 10% subject plasma (post-race); and 2) DMEM + 1% penicillin-streptomycin + 10% subject plasma (post-race) + TEMPOL (1 µM).

#### Statistical Analyses

Paired t-tests with a two-sided significance level of α = 0.05 were used to compare the change in ROS production, NO production, eNOS activation, total eNOS, aortic elastic modulus and cPWV and between pre- and post-race measures. Descriptive statistics including the mean and standard error are provided for each set of data. Simple linear regressions were used to determine the relation between central PWV and training history. Outliers were identified using ROUT Test for outliers (α = 0.05). All statistical calculations were conducted in GraphPad Prism 10.

### Multiomics Measurements

#### Metabolomics and Lipidomics Sample Preparation

Prior to metabolomics or lipidomics analysis, individual samples were placed in 1.5 ml tubes for either metabolite or lipid extraction. Twenty µL were suspended in 180 µL of water/methanol (50:50 v/v) for metabolite extraction or 180 µL of isopropanol/methanol (50:50 v/v) for lipid extraction. Suspensions were vortexed for 30 minutes at 4°C and then centrifuged for 10 minutes, 18,213g, 4°C. Supernatants were isolated for LC-MS.

#### Metabolomics and Lipidomics UHPLC-MS data acquisition and processing

Analyses were performed as previously published. ^27,28^ Briefly, the analytical platform employs a Vanquish UHPLC system (Thermo Fisher Scientific, San Jose, CA, USA) coupled online to a Q Exactive mass spectrometer (Thermo Fisher Scientific, San Jose, CA, USA). Polar metabolite extracts are resolved in singlicate over a Kinetex C18 column, 2.1 x 150 mm, 1.7 µm particle size (Phenomenex, Torrance, CA, USA) equipped with a guard column (SecurityGuard^TM^ Ultracartridge – UHPLC C18 for 2.1 mm ID Columns, Phenomenex, Torrance, CA, USA) using an aqueous phase (A) of water and 0.1% formic acid and a mobile phase (B) of acetonitrile and 0.1% formic acid for positive ion polarity mode, and an aqueous phase (A) of water:acetonitrile (95:5) with 1 mM ammonium acetate and a mobile phase (B) of acetonitrile:water (95:5) with 1 mM ammonium acetate for negative ion polarity mode. The column is equilibrated at 5% B, and upon injection of 10 μl of ex-tract, samples are eluted from the column using the solvent gradient: 0.5-1.1 min 5-95% B at 0.45 mL/min; hold at 95% B for 1.65 min at 0.45 mL/min, and then decrease to 5% over 0.25 min at 0.45 ml/min, followed by a re-equilibration hold at 5% B for 2 minutes at 0.45 ml/min. The Q Exactive mass spectrometer (Thermo Fisher Scientific, San Jose, CA, USA) is operated independently in positive or negative ion mode, scanning in Full MS mode (2 μscans) from 60 to 900 m/z at 70,000 resolution, with 4 kV spray voltage, 45 sheath gas, 15 auxiliary gas, AGC target = 3e6, maximum IT = 200 ms. Non-polar lipid extracts are re-solved over an ACQUITY HSS T3 column (2.1 x 150 mm, 1.8 µm particle size (Waters, MA, USA) using an aqueous phase (A) of 25% acetonitrile and 5 mM ammonium acetate and a mobile phase (B) of 90% isopropanol, 10% acetonitrile and 5 mM ammonium acetate. The column is equilibrated at 30% B, and upon injection of 10 μl of extract, samples are eluted from the column using the solvent gradient: 0-9 min 30-100% B at 0.325 mL/min; hold at 100% B for 3 min at 0.3 mL/min, and then decrease to 30% over 0.5 min at 0.4 ml/min, followed by a re-equilibration hold at 30% B for 2.5 minutes at 0.4 ml/min. The Q Exactive mass spectrometer (Thermo Fisher) is operated in positive ion mode, scanning in Full MS mode (2 μscans) from 150 to 1500 m/z at 70,000 resolution, with 4 kV spray voltage, 45 sheath gas, 15 auxiliary gas. When required, dd-MS2 is performed at 17,500 resolution, AGC target = 1e5, maximum IT = 50 ms, and stepped NCE of 25, 35 for positive mode, and 20, 24, and 28 for negative mode. Calibration is performed prior to analysis using the Pierce^TM^ Positive and Negative Ion Calibration Solutions (Thermo Fisher Scientific).

#### Metabolomics and Lipidomics Data analysis

Acquired data were converted from raw to mzXML file format using Mass Matrix (Cleveland, OH, USA). Samples were analyzed in randomized order with a technical mixture injected after every 10 samples to qualify instrument performance. Metabolite assignments are performed using accurate intact mass (sub-10 ppm), isotopic patterns, and retention time/spectral comparison to an in-house standard compound library (MSMLS, IROA Technologies, NJ, USA) using MAVEN (Princeton, NJ, USA). Lipidomics data are analyzed using LipidSearch 5.0 (Thermo Scientific), which provides lipid identification on the basis of accurate intact mass, isotopic pattern, and MS/MS fragmentation pattern to determine lipid class and acyl chain composition. Graphs, heat maps and statistical analyses (either T-Test or ANOVA), multivariate analyses including Principal Component Analysis (PCA), Partial Least Squares-Discriminant Analysis (PLS-DA), hierarchical clustering analysis (HCA), metabolite pathway enrichment analysis, and linear mixed models were prepared using MetaboAnalyst 5.0^29^

#### Proteomics sample preparation

Plasma samples were processed on S-Trap 96-well plates (Protifi, Huntington, NY) per manufacturer instructions. Briefly, ∼50 µg protein per well was solubilized in 5% SDS, reduced with 10 mM dithiothreitol (DTT) at 55 °C for 30 min, cooled to room temperature, and alkylated with 25 mM iodoacetamide (IAA) in the dark for 30 min. Phosphoric acid was added (final 2.5%), followed by six volumes of binding buffer (90% methanol, 100 mM triethylammonium bicarbonate [TEAB], pH 7.1). After gentle mixing, the protein solution was loaded onto S-Trap plates and centrifuged at 1,500 × g for 2 min; filters were washed 3× with 300 µL binding buffer. Sequencing-grade trypsin (Promega; 1 µg per well) in 125 µL digestion buffer (50 mM TEAB) was added, and plates were incubated at 37 °C for 6 h. Peptides were sequentially eluted with 100 µL each of (1) 50 mM TEAB, (2) 0.2% formic acid (FA) in water, and (3) 50% acetonitrile/0.2% FA. Eluates were pooled, lyophilized, and reconstituted in 0.1% FA (typically 500 µL) for LC-MS.

#### Proteomics data acquisition

For each sample, 20 µL of peptide solution was loaded onto individual Evotips (Evosep) for desalting and washed 3× with 200 µL 0.1% FA; 100 µL 0.1% FA was added to maintain hydration until analysis. Chromatography used a PepSep column (150 µm ID × 15 cm) packed with ReproSil-Pur C18 (1.9 µm, 120 Å) on an Evosep One system (30 samples/day preset, Evosep, Odense, Denmark). The LC was coupled via a CaptiveSpray nano-ESI source to a timsTOF Pro (Bruker Daltonics, Bremen, Germany) operated in DIA-PASEF mode.

The DIA-PASEF method comprised 32 dia-PASEF scans per cycle, each 100 ms, sampling the diagonal charge line for doubly/triply charged precursors with narrow 25 m/z precursor windows spanning m/z 100–1,700. A typical cycle time was 1.17 s (1 MS + 10 PASEF MS/MS scans). Low-abundance precursors with intensities >500 counts but below a target of 20,000 counts were repeatedly scheduled; dynamic exclusion was 0.4 min.

A project-specific spectral library was generated from pooled plasma/RBC digests fractionated at high pH and acquired by DDA-PASEF. For fractionation, pooled digest was separated on a Gemini-NH C18 column (4.6 × 250 mm, 3 µm) at 0.6 mL/min with mobile phase A: 20 mM ammonium bicarbonate (pH 10), and mobile phase B: 20 mM ammonium bicarbonate/75% acetonitrile (pH 10). The gradient was: 0–5% B (10 min), 5–50% B (80 min), 50–100% B (10 min), hold at 100% B (10 min). Ninety-six fractions were collected and concatenated to 24 (1+25+49+73, etc.), dried, reconstituted in 0.1% FA (80 µL), and 20 µL of each was loaded to Evotips for DDA-PASEF library acquisition.

#### Proteomics data processing and statistics

DIA raw files were processed in Spectronaut v17.6.230428.55965 (Biognosys). Processing used a project-specific library (“TEFR Base Mods”) with 23,756 targeted precursors and mutated decoys; fragment intensities were predicted by Spectronaut’s neural network. iRT calibration employed the Spectronaut iRT kit and Precision iRT with local non-linear regression; deamidated peptides were excluded from iRT calibration.

Identification thresholds were precursor q ≤ 0.01 (PEP ≤ 0.20) and protein q ≤ 0.01 at the experiment level (q ≤ 0.05 at run level), using dynamic decoy limiting (decoy/library fraction 0.1) and the kernel-density p-value estimator. Protein inference used the IDPicker workflow. Quantification was performed at MS1 (area) with automatic cross-run normalization enabled and interference correction on (minimum interferences: MS1 = 2, MS2 = 3). Grouping used protein group ID (major) and stripped peptide (minor) with TopN = 3 at both protein and peptide levels; proteotypicity filtering was disabled. Precursor filtering was “Identified (Qvalue)”; run-wise imputation was applied to missing values. Pathways were visualized using Cytoscape 3.10.4 (https://cytoscape.org).

## Results

### Multiomics analysis identifies significant associations between performance and vascular function mediators

Participant characteristics are presented in **Table 1**. Runners were followed by the research team along each stage in a trailer for a total of 64 stages and 4,486 kilometers from Bari, Italy to North Cape, Norway (**Figure 1A**). In addition to measuring central pulse wave velocity (cPWV) on site, isolated plasma was stored for animal and cell culture experiments, as well as multiomics characterization (**Figure 1B**). Proteomics, lipidomics, and metabolomics characterized 625, 643, and 237 plasma molecules, respectively (**Figure 1B, Supplementary Table 1**), which clearly distinguished samples taken before and after the race by partial least squares discriminant analysis (**Figure 1C**). There were a greater number of metabolites that changed between plasma sampled at baseline Pre- and Post-TEFR compared to proteins and lipids (**Figure 1D**). Notably, increased acylcarnitines indicated ongoing fatty acid oxidation throughout the race. This trend paralleled altered metabolites of arginine metabolism including creatine (increased) and guanidinoacetate (decreased). A similar alteration was observed with microbial metabolites that either increased (trimethylamineoxide [TMAO]) or decreased (indole-3-acetate, cresol), indicating an impact of the extreme ultra endurance race on the microbiome. Additionally, tyrosine metabolism was notably modulated (decreased – dityrosine, vanillate; increased – hippurate), while the co-factor tetrahydrobiopterin was the most significantly decreased metabolite.

**Figure 1.**
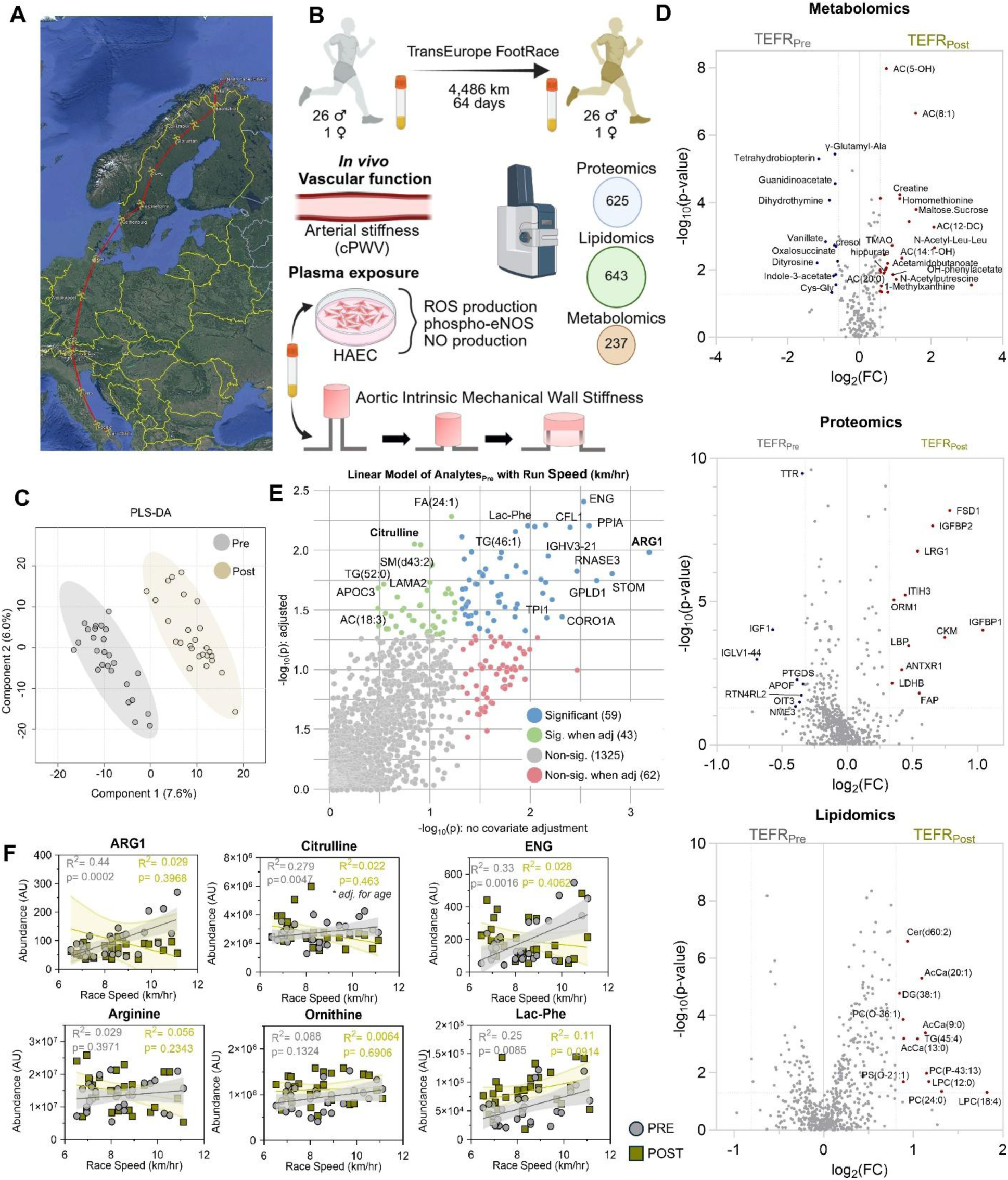
Multiomics and Vascular Function Analysis of the TransEurope Footrace. **(A)** Map of the 64-stage TransEurope Foot Race (TEFR) route spanning 4,486 km from Bari, Italy to North Cape, Norway. **(B)** Schematic of the study design. **(C)** Partial Least Squares Discriminant Analysis (PLS-DA) scores plot of multi-omics plasma data between Pre (silver) and Post (gold) timepoints. **(D)** Volcano plots of significant features in the metabolomics (top), proteomics (middle), and lipidomics (bottom) datasets. Significantly increasing and decreasing features are highlighted in red and blue, respectively. **(E)** Linear mixed model results of Pre-TEFR molecular correlates to subsequent average race speed. The x-axis represents significance without covariate adjustment, while the y-axis represents significance adjusted for age and sex. Blue dots indicate analytes significant in both models. **(F)** Representative scatter plots of top baseline molecular correlates with race speed, including Arginase 1 (ARG1), Citrulline, and soluble Endoglin (ENG). Silver circles represent Pre-race values; Gold squares represent Post-race values. R^2^ and p-values shown correspond to the Pre-race (silver) and Post-race (gold) correlations.

**Table 1:** Subject characteristics (mean ± SEM; N = 27)

|  | Male (n = 26) | Female (n = 1) |
| --- | --- | --- |
| **Age (years)** | 47.1 ± 10.7 | 45.0 |
| **BMI (kg/m<sup>2</sup>)** | 22.9 ± 2.1 | 22.4 |
| **SBP (mmHg)** | 130.2 ± 9.5 | 115 |
| **DBP (mmHg)** | 79.8 ± 6.9 | 75 |
| **Avg race time (hours)** | 536.3 ± 76.3 | 692.7 |
| **Avg race speed (km/hr)** | 8.6 ± 1.3 | 6.5 |

In support of alterations to the microbiome, lipopolysaccharide binding protein (LBP) emerged as one of the increased proteins, thereby suggesting damage to the intestinal endothelial barrier^30^. Also elevated were acute phase reactants orcomusoid-1 (ORM1), Leucine-rich alpha-2-glycoprotein 1 (LRG1), and downregulated transthyretin (TTR), indicative of ongoing inflammation. Notably, the decreased levels of IGF-1 in parallel to elevated IGFBP1 and IGFBP2 indicate ongoing metabolic dysfunction and shift towards catabolic rather than anabolic programs.

To determine which circulating molecules before the race correlated with participants’ subsequent average race speeds over the entirety of the TEFR, a linear mixed model correcting for age and sex was developed that identified 102 molecules (**Figure 1E**). Pathway enrichment revealed significant correlations with multiple proteins involved in hemostasis and platelet function, cell adhesion, cell motility and migration, stress and wound response, and regulation/morphogenesis/body fluids (**Supplementary Figure 1**). Among these significant variables were prominent features of arginine metabolism including Arginase 1 (ARG1) and citrulline, both of which were significantly correlated with race speed at baseline (R^2^ and p-value = 0.44, 0.0002 and 0.28, 0.0047, respectively) but not after the race (**Figure 1F**). Notably, citrulline was also correlated with runner age (**Supplementary Figure 1**), in line with previous findings from nearly 14,000 individuals^31^, but maintained significance after correction. Citrulline is either produced from arginine during nitric oxide (NO) production via nitric oxide synthase (NOS) or ornithine via ornithine transcarbamylase (OTC); however, these two substrates were not correlated with race speed. Other top predictive correlates with race speed included soluble endoglin (ENG), which is derived from vascular endothelial cells and is involved in angiogenesis^32^, and lactoyl phenylalanine (Lac-Phe), which is an appetite suppressant produced in response to exercise^33,34^ (**Figure 1F**). Collectively, these results point to a baseline homeostasis suggestive of NO production capacity and protection against inflammation, which ultimately associated with performance over the subsequent 64 days of running.

### Extreme ultrarunning elicits inflammation

In addition to identifying molecular markers at baseline that were predictive of performance, unsupervised hierarchical clustering of samples based on significantly changed proteins clearly separated time points, indicating a coordinated proteomic response (**Figure 2A**). Gene Ontology (GO) Molecular Function analysis highlighted a substantial inflammatory response related to the acute phase response and complement cascade (**Figure 2B**). Enriched cell component terms were dominated by extracellular region, blood microparticles, and secretory granule lumens, consistent with release of plasma proteins and vesicle cargo (**Figure 2C**). In addition, molecular function emphasized pattern-recognition and complement binding along with endoprotease inhibitor activity, reflecting the complement–protease network (**Figure 2D**). Representative proteins within the upregulated module included canonical acute-phase reactants (CRP, SAA4, ORM1, LBP, SERPINA1, LRG1). Within the complement cascade, the most significantly upregulated protein was the classical/lectin pathway component Complement (C4A), consistent with the clearance of sterile debris (**Figure 2E, Supplementary Figure 2**). In contrast, the rate limiting enzyme for the alternative pathway amplification loop, Complement Factor D (CFD), was significantly lower suggesting a compensatory dampening of this pathway. However, this regulation appeared compromised at the terminal effector stage. The pore-forming component C9 was significantly increased while the membrane-anchoring component C7 was decreased, suggestive of consumption via tissue deposition. In support, key fluid-phase regulators Clusterin (CLU) and Vitronectin (VTN) were significantly depleted (**Figure 2E**). These trends collectively indicate diminished sequestration of C5b-9 complexes as soluble MAC (sMAC) in favor of bioactive lytic pore assembly. Taken together, these data indicate that extreme ultrarunning elicits a coordinated innate immune program centered on sterile inflammation and terminal complement dysregulation.

**Figure 2.**
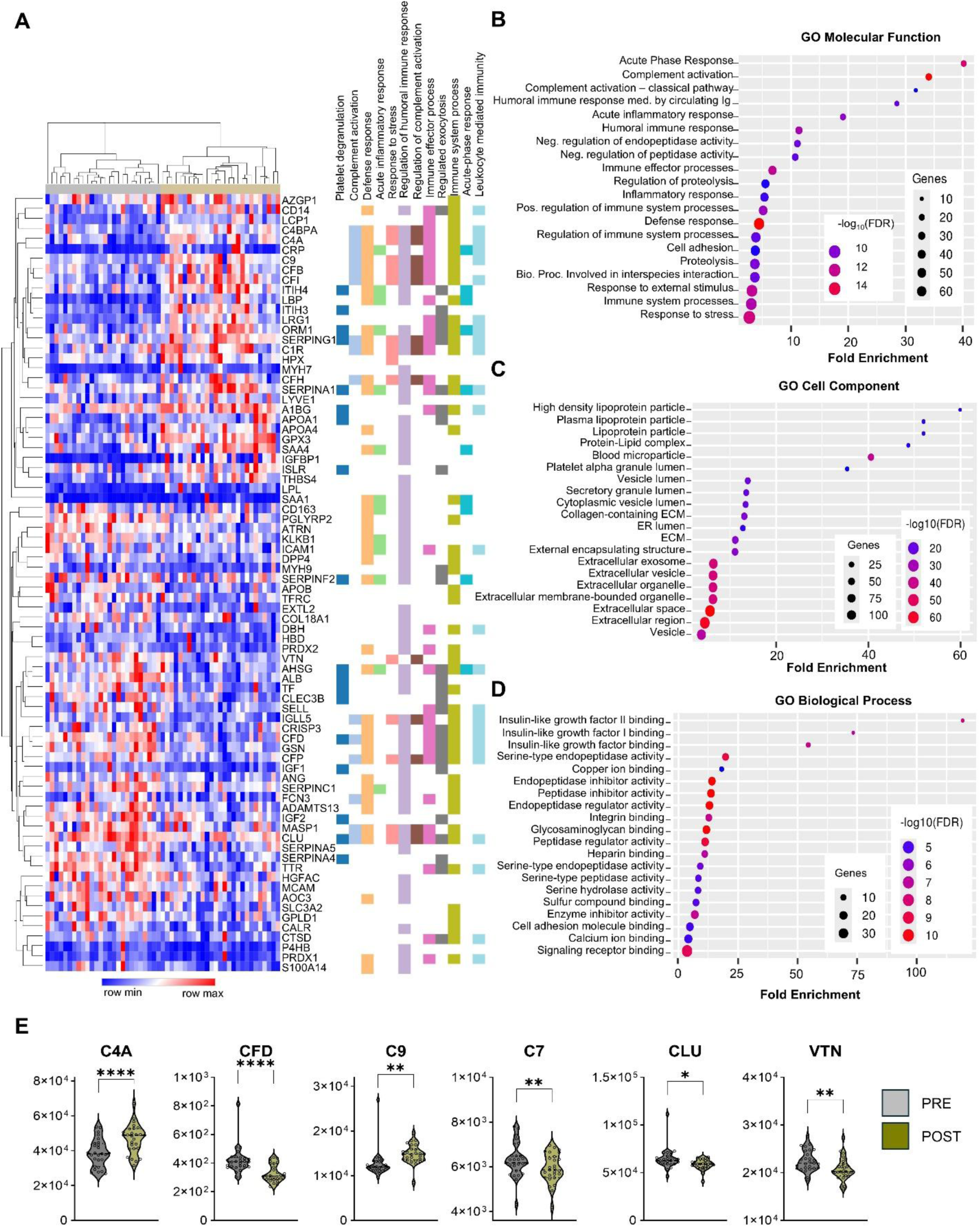
Extreme ultrarunning elicits a coordinated innate immune and protease-regulatory proteomic program. **(A)** Heatmap and unsupervised hierarchical clustering of significantly differentially abundant plasma proteins between Pre- and Post-TEFR timepoints. Rows represent individual proteins and columns represent participant samples. The color scale indicates relative abundance (red = high; blue = low). The right-hand annotation block identifies specific protein membership within key enriched functional clusters. **(B)** Gene Ontology (GO) enrichment analysis of significant proteins highlighting upregulation of biological processes related to inflammation. **(C)** GO Cellular Component analysis showing significant enrichment for proteins associated with lipoprotein particles, blood microparticles, and the extracellular space/vesicles. **(D)** GO enrichment analysis of molecular functions, emphasizing the regulation of proteolysis (serine-type endopeptidase/peptidase inhibitor activity) and insulin-like growth factor (IGF) binding. For B-D, dot size represents the number of genes associated with the term (count), color represents the statistical significance (-log₁₀(FDR)), and the x-axis represents the Fold Enrichment of the term.

### Race-induced lipid remodeling and perturbation of nitrogen metabolism

Class-level profiling showed broad lipidome shifts from pre- to post-race, dominated by increases in ceramides (Cer), acylcarnitines (AcCa), lysophosphatidylcholines (LPC), lysophosphatidylethanolamines (LPE), phosphatidylinositols (PI), lysophosphatidylinositols (LPI) and lysophosphatidic acids (LPA), with a concomitant decrease in phosphatidylglycerols (PG, **Figure 3A, B**). These trends were further evident at the species level. Increases were widespread across multiple LPC and ceramide species (**Figure 3B**). This expansion was driven most prominently by increases in very-long-chain species, including Cer(d18:1/24:0) and Cer(d18:1/24:1). In contrast, the level of the downstream metabolite sphingosine-1-phosphate (S1P) post-race was positively correlated with race speed (**Supplementary Figure 2**), highlighting a distinct decoupling between the accumulation of pathologic ceramides and the preservation of performance-associated S1P. Elevations in acylcarnitines across medium- to long-chain moieties, alongside a positive correlation between the CoA-precursor, pantothenate, and race speed (**Supplementary Figure 2**) illustrated the persistent demand for fat burning throughout the race that ultimately became limited by fatty acid oxidation capacity in the mitochondria. KEGG pathway enrichment analysis of significantly changed small molecule metabolites highlighted enrichment for arginine and proline metabolism, arginine biosynthesis, glycine/serine/threonine metabolism, and aromatic amino-acid biosynthesis (**Figure 3C**). Consistent with these pathway-level signals, targeted metabolites along the arginine–creatine axis were shifted post-race: creatine, ornithine, N-acetylputrescine, and N-acetylspermidine were higher, while guanidinoacetate, creatinine and proline were lower (**Figure 3D**). At steady state, these relative levels highlight increased flux through creatine and polyamine biosynthesis for tissue regeneration rather than NO production via eNOS. In support, the ECM-stabilizing protein ITIH3, as well as ECM-degrader Fibroblast Activator Protein (FAP), and ECM-remodeling protein ANTXR1 were all significantly elevated post-race (**Supplementary Figure 1**).

**Figure 3.**
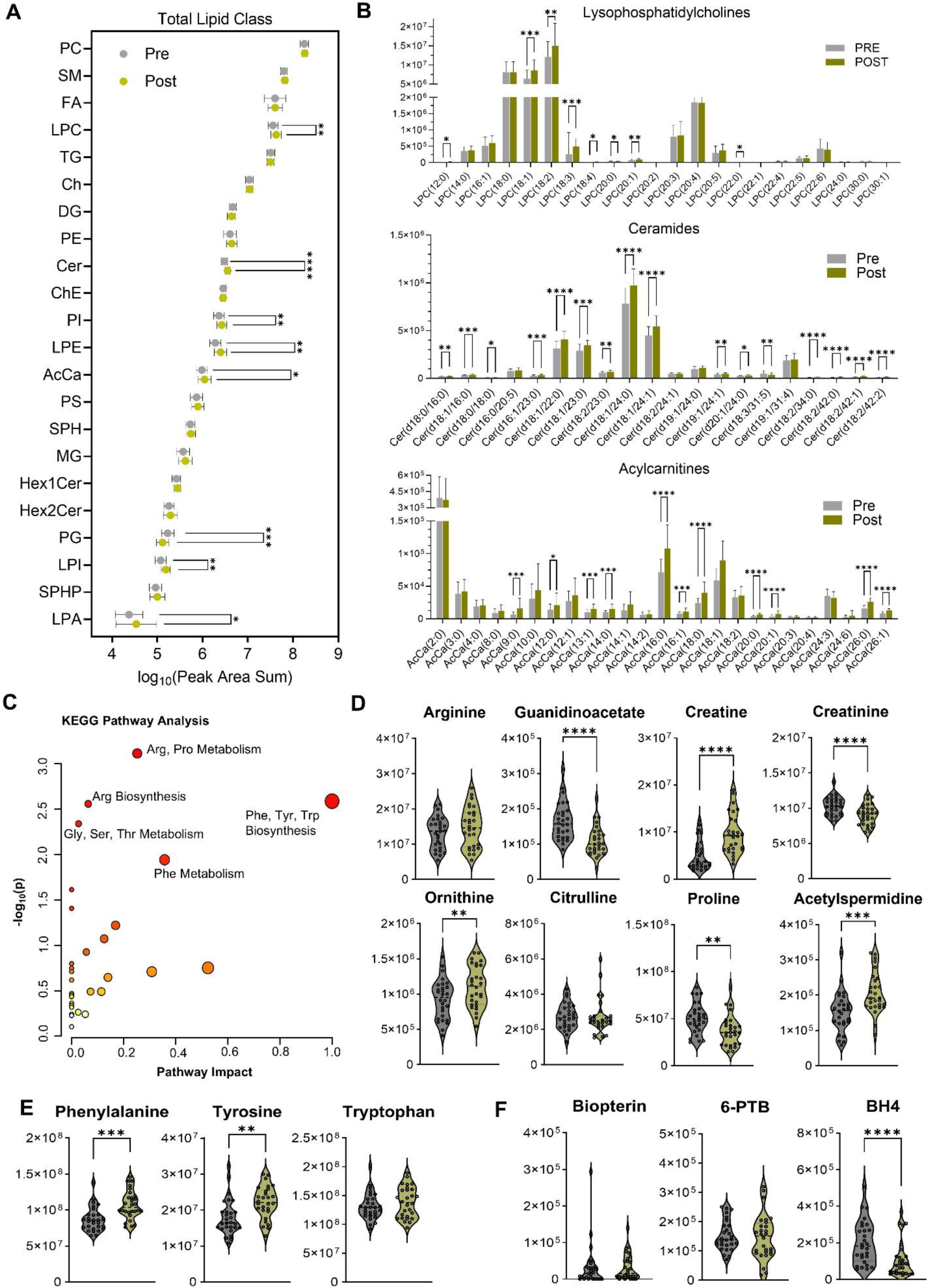
Systemic remodeling of the plasma lipidome and nitrogen metabolism following the TEFR. **(A)** Changes in total abundance of lipid classes in plasma collected Pre- (silver) and Post- (gold) TEFR. Classes include Phosphatidylcholines (PC), Lysophosphatidylcholines (LPC), Ceramides (Cer), Acylcarnitines (AcCa), and Phosphatidylglycerols (PG). Data are presented as mean ± SEM of the log_10_ sum of peak areas for each class. **(B)** Relative abundance of specific lipid species belonging to LPC, Cer, and AcCa classes. **(C)** KEGG pathway enrichment analysis of significantly altered small molecule metabolites. Node color is based on pathway significance, which is also denoted on the y-axis. **(D)** Violin plots showing the relative abundance of metabolites associated with the Arginine-Creatine axis (Arginine, Guanidinoacetate, Creatine, Creatinine) and polyamine synthesis (Ornithine, Acetylspermidine). **(E)** Relative abundance of Aromatic Amino Acids (Phenylalanine, Tyrosine, Tryptophan). **(F)** Relative abundance of pterin metabolites, including Biopterin, 6-Pyruvoyltetrahydropterin (6-PTB), and tetrahydrobiopterin (BH4). For D-F, violin plots display the distribution of individual data points. *P < 0.05, **P < 0.01, ***P < 0.001, ****P < 0.0001 by paired t-test.

### Post-race plasma increases arterial stiffness via ROS without altering endothelial NO

Given that the multiomics analysis suggested a systemic stress phenotype, we first assessed plasma-mediated oxidative stress in a vascular cell culture model. Consistent with the circulating molecular profile of plasma, ROS production in HAECs was significantly elevated after post-TEFR plasma exposure **(Figure 4A)**. Because oxidative stress often impairs vascular function via reduced nitric oxide (NO) bioavailability, we subsequently assessed NO production and eNOS activation. Surprisingly, NO production in HAECs did not change after exposure to pre- versus post-TEFR plasma **(Figure 4B)**. Furthermore, neither eNOS phosphorylation (at the Ser1177 or Thr495 sites) nor total eNOS abundance were significantly altered **(Figure 4C-E)**. Despite the preservation of NO synthesis parameters *ex vivo*, we next sought to determine if the post-race plasma could directly induce higher aortic elastic modulus, considering oxidative stress can independently drive aortic stiffening. We found that elastic modulus of thoracic aorta rings isolated from healthy intervention naïve mice was significantly increased following exposure to participants’ post-race plasma relative to exposure to the same participants’ pre-race plasma **(Figure 4F)**. To determine if this stiffening was mechanistically linked to the observed oxidative stress, we repeated the assay in the presence of the membrane-permeable superoxide dismutase mimetic, TEMPOL. Co-incubation with TEMPOL significantly lowered post-race plasma-induced elastic modulus **(Figure 4G)**, confirming a ROS-dependent component of the stiffening response induced by the systemic circulation after the TEFR. Finally, complementary to these serum-induced vascular changes, cPWV measured *in vivo* was significantly elevated after the race **(Figure 4H)**, indicating that the circulating factors identified here contribute to increased arterial stiffness *in vivo* in the athletes participating in the TEFR.

**Figure 4.**
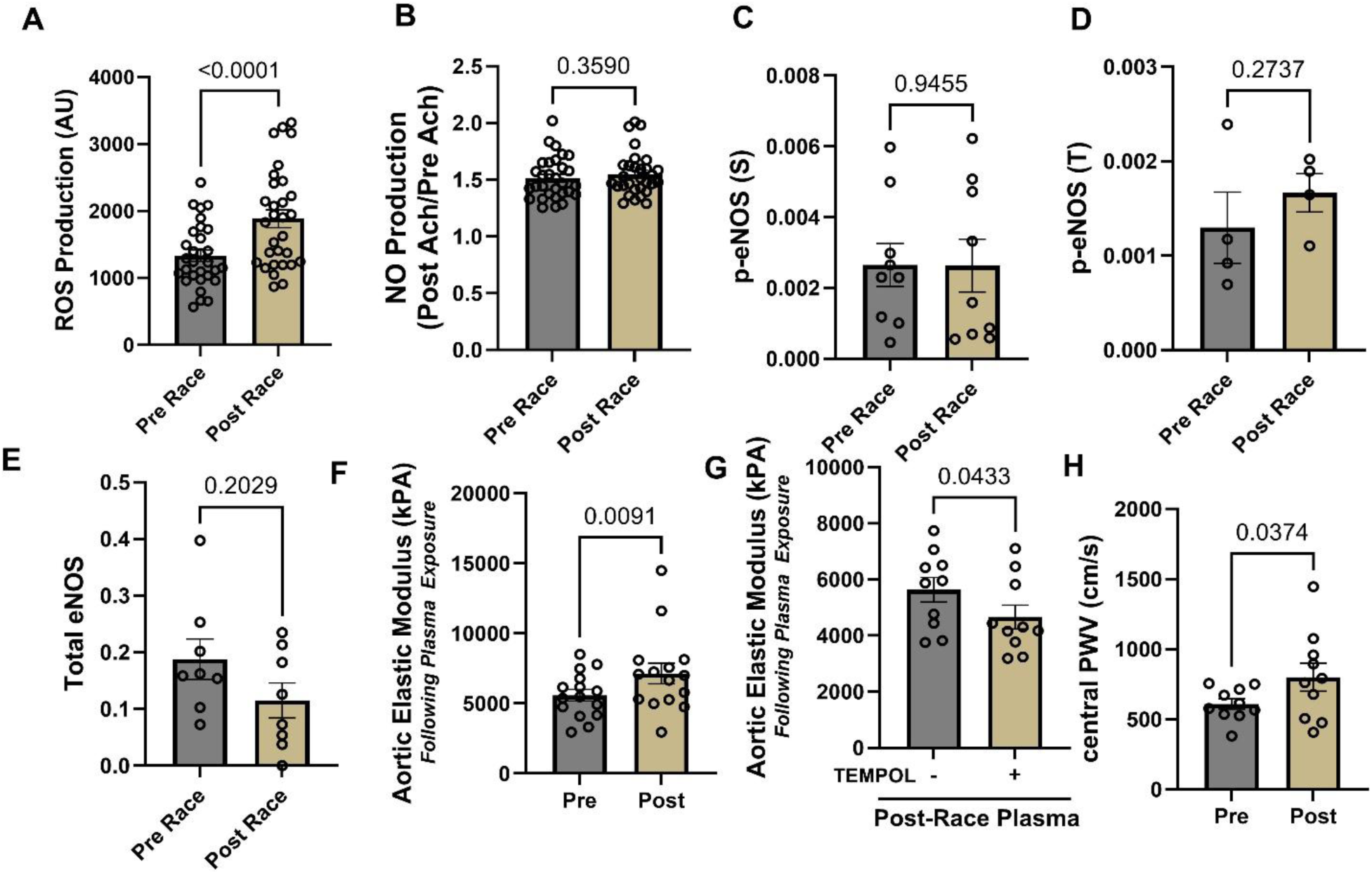
Impact of Ultrarunning on Vascular Function. **(A)** Quantification of reactive oxygen species (ROS) production in Human Aortic Endothelial Cells (HAECs) following incubation with plasma collected Pre- and Post-TEFR. **(B)** Quantification of nitric oxide (NO) production in HAECs exposed to Pre- and Post-TEFR plasma. Densitometric analysis of eNOS phosphorylation at **(C)** Ser1177 and **(D)** Thr495, and **(E)** total eNOS protein abundance in HAECs exposed to participant plasma. **(F)** Ex vivo aortic elastic modulus (stiffness) of naïve murine thoracic aortic rings following incubation with Pre- and Post-TEFR plasma. **(G)** Ex vivo aortic elastic modulus of murine aortic rings incubated with Post-TEFR plasma in the presence or absence of the superoxide dismutase mimetic TEMPOL. **(H)** In vivo central pulse wave velocity (cPWV) measured in TEFR participants before (Pre) and after (Post) the 4,500-km race. Data are presented as mean ± SEM. Significance determined by paired t-test (A-F, H) or paired t-test comparing vehicle vs. TEMPOL (G).

### Molecular correlates of *ex vivo* NO, ROS, and *in vivo* cPWV

Given the significant alterations in vascular function observed after the TEFR, we next sought to determine which molecules correlate with select parameters to better define the circulating milieu to which vascular cells are exposed. Several molecules showed correlations with cPWV (**Figure 5A**). Caffeine and its metabolites theobromine and 1-methylxanthine were top positive correlates (**Figure 5, Supplementary Figure 3**), reproducing established relations^35,36^ and providing validity to this approach. Additional correlates included sugar alcohols (arabitol/xylitol/ribitol which are not independently resolved), dodecanoic acid (FA[12:0]), alanine aminotransferase (GPT), fibronectin type III (FSD1), complement C4A, eicosasphinganine (SPH[d20:0]), β-hydroxybutyrate, and chymotrypsinogen B1/B2 (CTRB1/2). While plasma-induced NO production itself was not significantly impacted (**Figure 4B**), plasma-analyte correlations with endothelial NO output (**Figure 5B**) revealed positive correlation with Complement C8 gamma (C8G), and α-1-acid glycoprotein (ORM1), while top negative correlates included eNOS inhibitor dimethylarginine^37^, Complement C7 and prekallikrein (KLKB1). Finally, correlation between multi-omics data and corresponding HAEC ROS production after plasma exposure (**Figure 5C**) revealed prominent inverse association with the serine protease inhibitor SERPINA5 (r = -0.58, p = 5.9×10⁻^6^), which was significantly decreased after the TEFR (**Supplementary Figure 2**) – and positive associations with Complement 4 binding protein alpha (C4BPA) and lactate dehydrogenase B (LDHB), and Complement C9. Taken together these results implicate complement/coagulation proteins and proteases as major correlates of endothelial NO production and oxidative stress *ex vivo*, while C4A, sphingolipids, energy-stress metabolites (ketone bodies) link to cPWV *in vivo*.

**Figure 5.**
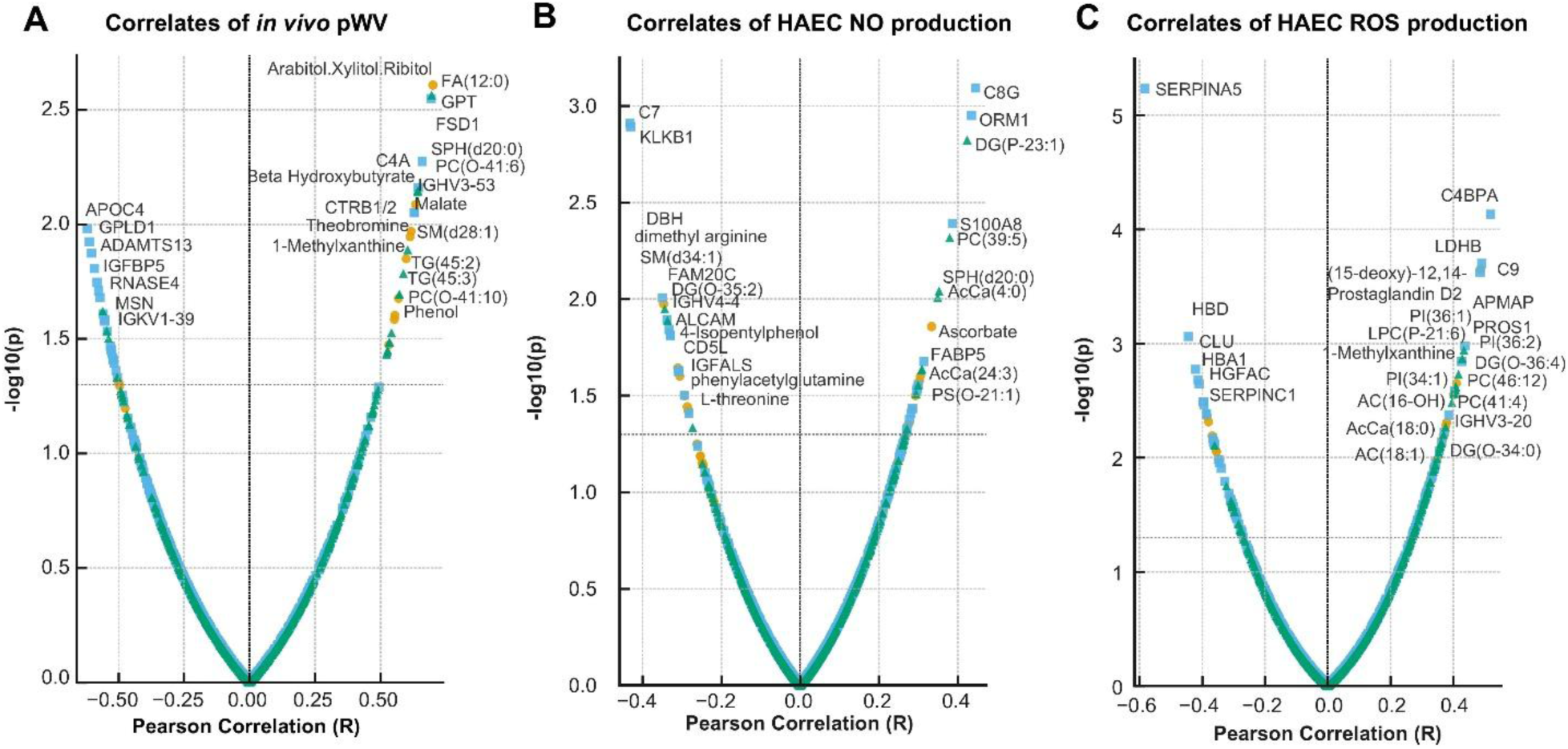
Multiomics Correlations with Vascular Function. Pearson correlations and p-values are presented for **(A)** pulse wave velocity, **(B)** HAEC nitric oxide (NO) production after plasma exposure, and **(C)** HAEC reactive oxygen species (ROS) production after plasma exposure. The top 20 correlates for each parameter are labeled.

## Discussion

To elucidate the molecular determinants of impaired vascular function following an extreme duration multi-stage endurance race, we integrated plasma metabolomics, lipidomics, and proteomics with vascular phenotyping from runners completing the 64-stage, 4,486-km TEFR. Our findings demonstrate that extreme ultra-endurance running sustained daily for two months profoundly alters the systemic molecular milieu, resulting in an oxidative associated with vascular dysfunction. We observed that post-race plasma increased endothelial ROS production without altering measured endothelial NO production. This plasma increased stiffness in murine aortic rings *ex vivo*, mirroring the increased cPWV observed in the runners. Multiomics analysis provided insights into the physiological impact of this extreme endurance event as well as how the circulating molecular milieu is associated with vascular function. Specifically, we identified coordinated remodeling of amino acid metabolism, evidence of incomplete fatty acid oxidation with bioactive lipid accumulation, and robust activation of inflammatory programs including the acute phase response, complement, and coagulation pathways.

Given the requisite training status to complete an event of this length, and in conjunction with the variation in running duration (the range of running time for participants included in this study was nearly 286 hours), we hypothesized that baseline circulating molecules might reflect pathways that confer performance resilience while post-race alterations would reveal mechanisms of physiological stress. At baseline before the start of the TEFR, ARG1 emerged as a top positive correlate of average run speed over the subsequent 2 months. As a urea cycle enzyme, ARG1 competes with eNOS for the shared substrate, arginine. In addition, the eNOS product citrulline also positively correlated with average run speed thus suggesting that baseline NO-related vascular biology may influence performance capacity during the course of the TEFR. However, the race induced significant perturbations in this pathway. A rerouted metabolic signature towards creatine synthesis may reflect modified energetic demands throughout the race. In addition, elevated ornithine alongside increased N-acetylputrescine and N-acetylspermidine indicate upregulation of polyamine synthesis to sustain tissue repair^38^. Moreover, an offshoot of this pathway also generates proline, which may be rerouted towards collagen synthesis for wound healing as has been observed in previous TEFR participants^39^.

Significant elevations in phenylalanine and tyrosine suggest either sustained proteolysis or decreased metabolic flux through aromatic amino acid pathways, or a combination thereof. Metabolism of these aromatic amino acids requires the cofactor BH4 as does eNOS. Decreased circulating levels of BH4 by the end of the race thus suggest decreased systemic availability of this cofactor. In support, several tyrosine catabolites were significantly downregulated after the race. Under conditions of cofactor deficiency, eNOS produces superoxide rather than NO, contributing to an increasingly oxidative environment^40^. Indeed, eNOS protein levels were not significantly different while ROS production was significantly elevated. These results thus suggest that the vascular compromise of extreme endurance is not likely driven by arginine exhaustion, but by outweighed demand of alternative metabolic circuits potentially mediated by diminished BH4 availability and eNOS uncoupling.

Beyond the arginine-biopterin axis, the simultaneous elevation of ceramides and acylcarnitines highlight a pro-inflammatory bioenergetic stress. The accumulation of medium- and long-chain acylcarnitines is consistent with incomplete fatty acid oxidation, in which fatty acid mobilization exceeds downstream oxidative capacity. This metabolic profile mirrors the accumulation of acylcarnitines observed after heart failure^41^, in patients suffering from post-acute sequalae of COVID-19^42^, or in elite cyclists after reaching volitional exhaustion^43^. In this cohort, the extent of this bioenergetic strain may contribute to the observed accumulation of downstream ceramides, particularly the very long chain species Cer(18:1/24:1), which is an established circulating biomarker for loss of muscle mass^44^. In addition, ceramides can impair vascular function by activating protein phosphatase 2A (PP2A), which dephosphorylates and uncouples eNOS^45^. These stresses collectively result in superoxide generation, leading to oxidative stress.

Excessive ROS production triggers damage-associated molecular patterns (DAMPs), initiating a systemic immune response. Consistent with reports on ultra endurance-induced proteomic inflammatory signatures^46,47^, our proteomic analysis here revealed a significant dysregulation of the complement system and coagulation pathways. The post-race plasma profile was dominated by acute-phase reactants and complement activation. The specific elevation of the terminal complement pathway (C7, C8, C9) is of particular interest. Unlike upstream activation, the assembly of the sub-lytic Membrane Attack Complex (MAC) on endothelial surfaces can induce calcium influx and P-selectin expression, thereby promoting a pro-thrombotic state^48^. We identified specific molecular correlates linking this inflammatory milieu to both endothelial ROS and NO production. The strongest negative correlate with endothelial cell ROS production was SERPINA5, a member of the serine protease inhibitor (SERPIN) family of proteins^49^. Notably, SERPINA5 contributes to the degradation of urokinase plasminogen activator^50^, which itself has been shown to induce endothelial ROS production^51^. Thus, lower SERPINA5 may reflect loss of a protective protease-regulatory signal associated with greater oxidative stress. Conversely, we observed negative or positive correlations, respectively, between NO production and C7 and C8G. C7 is a key component in the terminal pathway of complement activation and essential for the assembly of the terminal C complex (TCC) that binds to the EC surface to trigger activation and inflammation^52^ that can impair NO production. This process is neutralized in part by surface CD59^53^. Interestingly, the pre-TEFR levels of CD59 were also a top positive correlate with average running speed throughout the race thus suggesting that a basal ability to inhibit soluble C7 may promote endurance capacity. On the other hand, C8G, which is also involved in TCC formation, has been established as a factor in maintaining blood brain barrier integrity and reducing pro-inflammatory signaling by blocking the sphingosine 1 phosphate receptor 2 (S1PR2) on endothelial cells^54,55^. Notably, post-TEFR levels of S1P itself were positively correlated with running speed. C8G could thus serve as a compensatory mechanism to cope with exercise-elicited production of S1P in endothelial cells responding to laminar shear stress^56^, or in red blood cells^57^ to modulate oxygen offloading kinetics^58^. S1P synthesis also relies on ceramide catabolism and may be upregulated, in part, to protect against lipotoxic accumulation.

The complement profile is consistent with a state of regulated sterile inflammation. The immune response was dominated by the Classical and Lectin pathway activation with specific suppression of the Alternative Pathway, thus indicating a compensatory mechanism to limit systemic toxicity while maintaining debris clearance. Relative factor abundances (increased C9, decreased C7), together with reduced VTN and CLU^59^, are compatible with altered terminal complement dynamics and raise the possibility of increased endothelial exposure to bioactive terminal complement complexes. The depletion of CLU observed here stands in stark contrast to the physiological response to moderate exercise observed by De Miguel et al., who identified CLU as an anti-inflammatory factor upregulated by voluntary running that binds cerebral endothelial cells and inhibits complement activation^60^. CLU elevations in that study were observed after shorter interval and moderate training, which differs from the immense exercise volume of the TEFR. These results raise the possibility that, beyond a certain exercise burden, protective anti-inflammatory responses become insufficient to fully buffer sterile inflammatory stress.

A key finding of this study is the dissociation between endothelial cell NO production and ROS-mediated large artery stiffening induced by the circulating milieu. In endothelial cells exposed to post-race plasma, we observed no significant change in NO production, eNOS protein levels, or phosphorylation status. However, ROS production was significantly elevated. This suggests that the TEFR plasma induces vascular dysfunction by overwhelming the redox balance rather than silencing NO synthesis. In support, post-race plasma acutely increased the stiffness of naïve murine aortic rings, which was attenuated when co-incubated with the superoxide dismutase mimetic TEMPOL. While physiological ROS signaling is required for adaptation and vasodilation in skeletal muscle^61,62^, excessive oxidative stress damages structural proteins and scavenges NO, reducing its bioavailability despite preserved synthesis. Thus, the large artery stiffening observed here could be driven by a ROS-dependent reduction in NO bioavailability and direct oxidative damage to the vessel wall.

The existing literature on vascular function in ultra-endurance athletes is complex. The acute response of large artery function to an ultra-marathon is highly variable and appears dependent on distance and duration: a study of the 161-km Western States Ultramarathon reported no immediate change^63^, while others observe an acute increase in stiffening, particularly following longer distances^64,65^. Peripheral vascular function, including brachial artery endothelial function, tends to be either preserved or experiences a transient decrease that recovers within 24 hours^66,67^, suggesting that the microvasculature is generally more resilient or recovers faster than the central elastic arteries. Moreover, runners completing the 330-km TOR des Géants exhibited markers of cardiac fibrosis, ischemia, and inflammation that persisted for days^68^, in addition to World Mountain Running Championship participants who have shown pro-thrombotic shifts^69^ that mirror complement pathway activation observed here.

The dependency of vascular damage on duration and cumulative load raises a fundamental question regarding the threshold between hormetic adaptation and pathologic injury. While moderate exercise bolsters antioxidant defenses, the dose of exercise endured during the TEFR appears to overwhelm these compensatory mechanisms. The proteomic overlap seen here and in the UTMB mirrors multiomics signatures observed in patients who have suffered endotheliopathic, hemorrhagic, and traumatic injuries^70^, indicating that at this volume, exercise ceases to be a signaling stimulus and becomes a systemic insult. Our data imply that the human body’s capacity to adapt to acute ultra-endurance efforts is robust, but the chronic repetition of these events without adequate recovery drives the systemic molecular milieu toward a maladaptive profile. It is thus interesting to consider the impact that competing in multi-stage endurance races or multiple individual events over a relatively short period of time (e.g. multiple ultramarathons over the course of a year) have on the epigenetic landscape as well as microbiome species diversity. Indeed, Lac-Phe that was observed here to associate with performance capacity is a microbial metabolite (in addition to p-cresol^71^, indole-3-acetate^72^, and TMAO that is associated with cardiovascular pathology^73^ in part due to its role in arterial stiffening^74^), which is produced by the condensation of lactate and phenylalanine^75^ in response to exercise^33,34^. Moreover, lactate is a substrate for histone modification^76^. Considering the exertion-dependent accumulation of lactate and other post-translational modification substrates including succinate^77^, future studies should examine the impact of repeat exertive bouts with limited recovery intervals on longer term adaptations (both beneficial and deleterious) to exercise.

This study has limitations inherent to the extreme nature of the event. First, logistical hurdles precluded collection of multiple longitudinal samples in this cohort, which would have provided better temporal resolution of the altered molecular physiology we observed here. Due to limited sample volumes, *in vivo* endothelial function could not be assessed to validate *ex vivo* cell culture-based findings. In addition, the cPWV sample size was relatively small, though we did observe mechanistic corroboration in murine tissues. Finally, due to limitations with study recruitment, this cohort was nearly entirely male. In consideration of sex-based differences in exercise responses^78^, as well as the complement system^79^, follow up studies must be more balanced.

Collectively, this study is the first to link the systemic multiomics signature of extreme ultrarunning directly to key vascular function phenotypes. We demonstrate that the circulating milieu following chronic ultra-endurance running promotes a systemic inflammatory and oxidative state. Furthermore, our *ex vivo* findings suggest that this plasma-induced vascular burden is driven primarily by excessive oxidative stress rather than a direct failure of endothelial NO synthesis. This phenotype is characterized by incomplete fatty acid oxidation, accumulation of proinflammatory lipids and proteins, complement activation, and modified amino acid metabolism associated with depletion of BH4. Future work should characterize the temporal resolution of these perturbations. Longitudinal follow-up of these unique athletes could determine if these acute functional changes translate to long-term vascular remodeling. Indeed, despite the deleterious impacts we observed after running for 4,486-km, routinely active ultra-endurance runners often exhibit a beneficial chronic adaptation, showing lower baseline central arterial stiffness compared to age-matched controls^80,81^. These data nominate the complement-redox axis, ceramide metabolism, and BH4-related pathways as candidate targets for future mechanistic and interventional studies..

## Acknowledgments

The authors would like to acknowledge Daniel H Craighead for his advice, technical insights, and constructive feedback during the execution of this study.

## Funding

This work was supported by funds from the National Institutes of Health (NIH) T32 GM159536 (CMC), T32 HL007171 (FC), NIH, National Heart, Lung, and Blood Institute (NHLBI) awards R01HL146442, R01HL149714, R01HL148151 (AD), NHLBI R00HL159241 (ZSC), NIH U01AR071124 (ZSC), National Cancer Institute award R01CA292482 (TN), and National Institute of Aging award U54AG062319 (TN).

## Author contributions

Conceptualization: ZSC, TN, DRS, AST, KRL

Methodology: CMC, CCU, FC, MD, SPF, KRL, DHC, UHS, ZSC

Investigation: CMC, CCU, KRL, TN

Visualization: TN, KRL, ZSC

Supervision: TN, ZSC, KRL

Writing – original draft: TN, CMC, ZSC, KRL

Writing – review & editing: All authors

## Competing interests

Although unrelated to the contents of this manuscript, AD and TN are founders of Omix Technologies Inc. and scientific advisory board members for Hemanext Inc. AD is a scientific advisory board member for Macopharma Inc. The remaining authors declare no competing financial interests.

## Data and materials availability

All data are available in the main text or the supplementary materials.

---

**Supplementary Figure 1.**
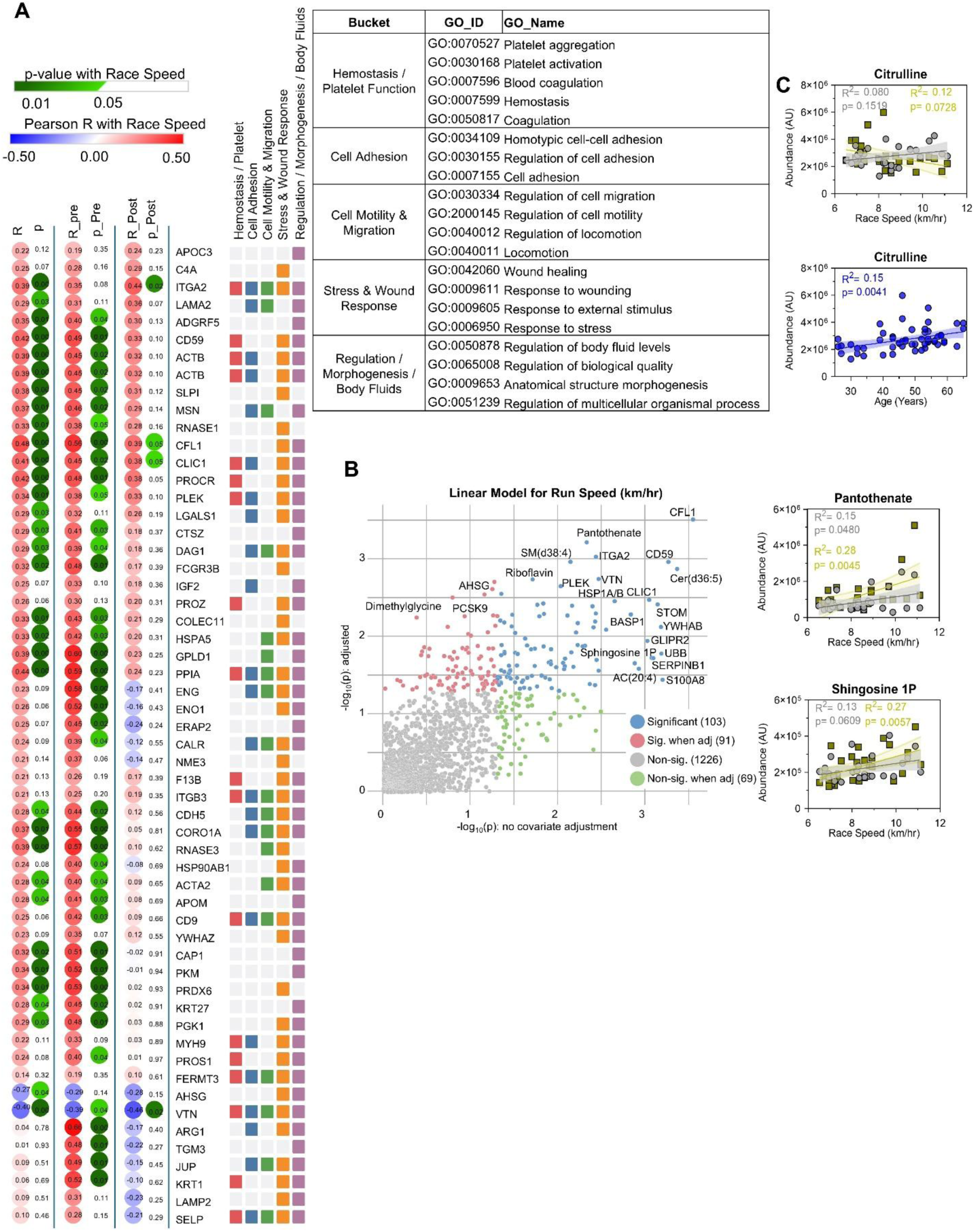
(A) Principal Component Analysis of plasma samples from Trans European Footrace participants before (gray) and after (gold) the race. (B) A linear mixed model for Run Speed using multiomics analytes at Pre and Post time points. (C) A correlation plot for citrulline without correction for participant age (top) and a plot demonstrating Pearson correlation between citrulline levels and age (bottom).

**Supplementary Figure 2.**
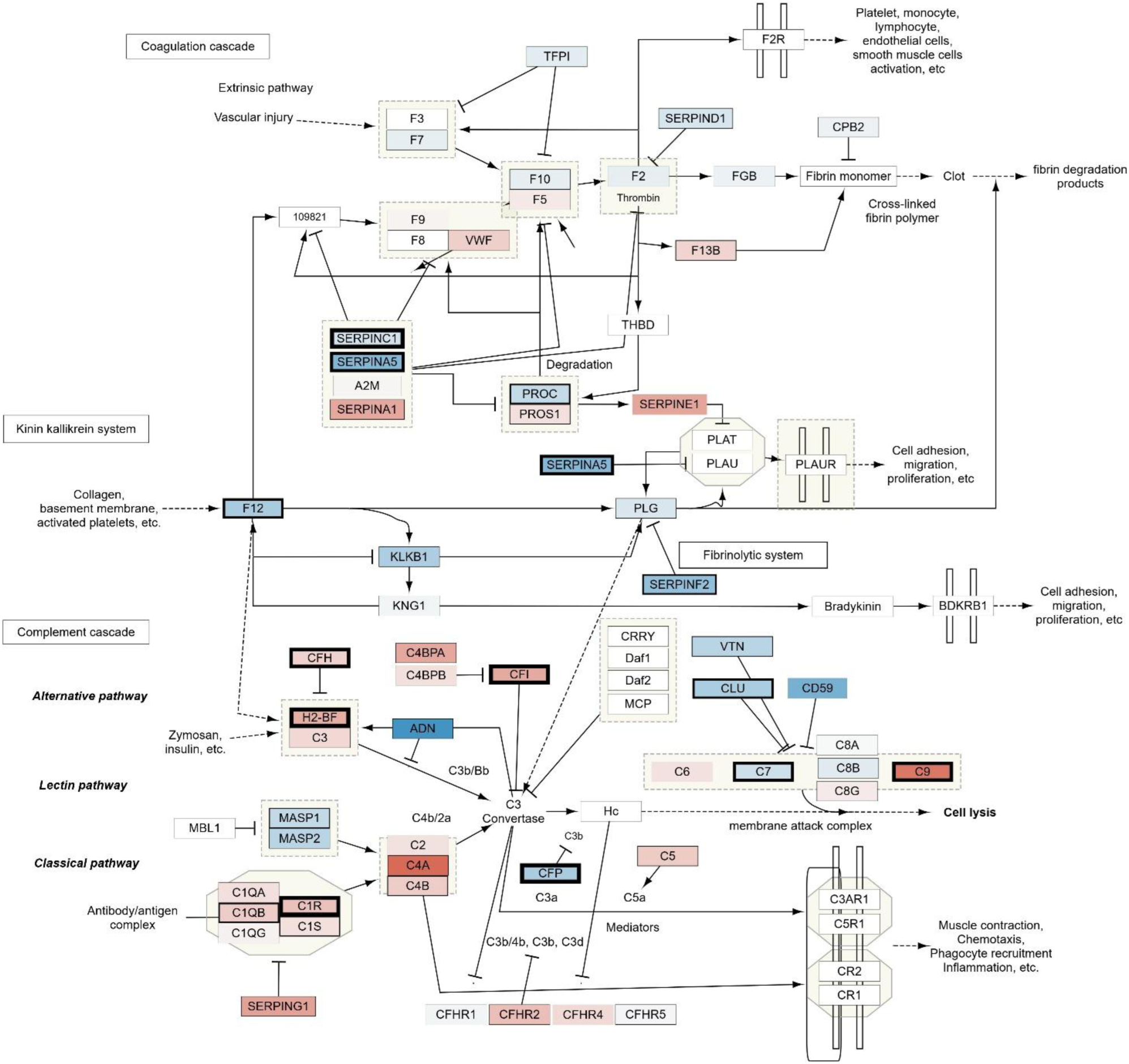
Coagulation and Complement Cascades. Network visualization (WikiPathways) is overlaid with proteomic abundance data comparing Post to Pre. Node fill color indicates the magnitude of change (Log2 Fold Change), with red representing upregulation and blue representing downregulation relative to controls. Statistical significance is visualized via node border weight.

**Supplementary Figure 3.**
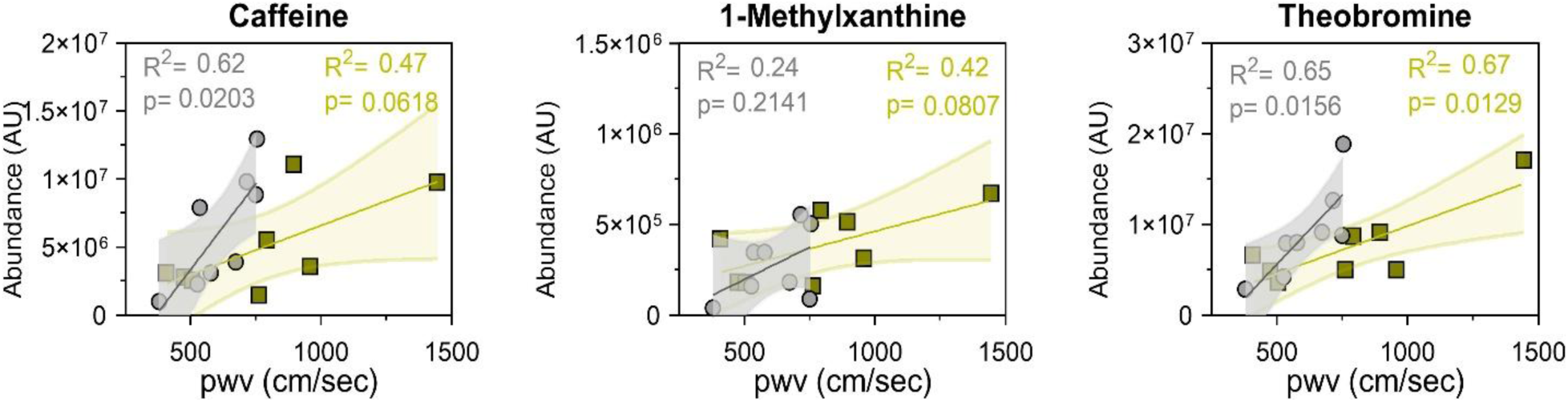
Caffeine and metabolites correlate with pulse wave velocity (pWV).

## Notes

### Competing Interest Statement

The authors have declared no competing interest.

## References

1. Ungvari Z, Tarantini S, Donato AJ, Galvan V, Csiszar A. Mechanisms of vascular aging. Circulation research. 2018;123:849–867.

2. Moreira JBN, Wohlwend M, Wisløff U. Exercise and cardiac health: physiological and molecular insights. Nat Metab. 2020;2:829–839. doi: 10.1038/s42255-020-0262-1

3. Lear SA, Hu W, Rangarajan S, Gasevic D, Leong D, Iqbal R, Casanova A, Swaminathan S, Anjana RM, Kumar R, et al. The effect of physical activity on mortality and cardiovascular disease in 130 000 people from 17 high-income, middle-income, and low-income countries: the PURE study. Lancet. 2017;390:2643–2654. doi: 10.1016/s0140-6736(17)31634-3

4. Seals DR, Nagy EE, Moreau KL. Aerobic exercise training and vascular function with ageing in healthy men and women. J Physiol. 2019;597:4901–4914. doi: 10.1113/jp277764

5. Taddei S, Galetta F, Virdis A, Ghiadoni L, Salvetti G, Franzoni F, Giusti C, Salvetti A. Physical activity prevents age-related impairment in nitric oxide availability in elderly athletes. Circulation. 2000;101:2896–2901. doi: 10.1161/01.cir.101.25.2896

6. Eskurza I, Monahan KD, Robinson JA, Seals DR. Effect of acute and chronic ascorbic acid on flow-mediated dilatation with sedentary and physically active human ageing. J Physiol. 2004;556:315–324. doi: 10.1113/jphysiol.2003.057042

7. Vaitkevicius PV, Fleg JL, Engel JH, O’Connor FC, Wright JG, Lakatta LE, Yin FC, Lakatta EG. Effects of age and aerobic capacity on arterial stiffness in healthy adults. Circulation. 1993;88:1456–1462. doi: 10.1161/01.cir.88.4.1456

8. Tanaka H, Dinenno FA, Monahan KD, Clevenger CM, DeSouza CA, Seals DR. Aging, habitual exercise, and dynamic arterial compliance. Circulation. 2000;102:1270–1275. doi: 10.1161/01.cir.102.11.1270

9. Tanaka H, DeSouza CA, Seals DR. Absence of age-related increase in central arterial stiffness in physically active women. Arterioscler Thromb Vasc Biol. 1998;18:127–132. doi: 10.1161/01.atv.18.1.127

10. Moreau KL, Donato AJ, Seals DR, DeSouza CA, Tanaka H. Regular exercise, hormone replacement therapy and the age-related decline in carotid arterial compliance in healthy women. Cardiovasc Res. 2003;57:861–868. doi: 10.1016/s0008-6363(02)00777-0

11. DeSouza CA, Shapiro LF, Clevenger CM, Dinenno FA, Monahan KD, Tanaka H, Seals DR. Regular aerobic exercise prevents and restores age-related declines in endothelium-dependent vasodilation in healthy men. Circulation. 2000;102:1351–1357. doi: 10.1161/01.cir.102.12.1351

12. Pierce GL, Eskurza I, Walker AE, Fay TN, Seals DR. Sex-specific effects of habitual aerobic exercise on brachial artery flow-mediated dilation in middle-aged and older adults. Clin Sci (Lond). 2011;120:13–23. doi: 10.1042/cs20100174

13. Ho LYW, Kwan RYC, Yuen KM, Leung WC, Tam PN, Tsim NM, Ng SSM. The Effect of Aerobic Exercises on Arterial Stiffness in Older People: A Systematic Review and Meta-Analysis. The Gerontologist. 2023;64. doi: 10.1093/geront/gnad123

14. Eijsvogels TMH, Thompson PD, Franklin BA. The “Extreme Exercise Hypothesis“: Recent Findings and Cardiovascular Health Implications. Curr Treat Options Cardiovasc Med. 2018;20:84. doi: 10.1007/s11936-018-0674-3

15. Mastaloudis A, Leonard SW, Traber MG. Oxidative stress in athletes during extreme endurance exercise. Free Radic Biol Med. 2001;31:911–922. doi: 10.1016/s0891-5849(01)00667-0

16. Oh HS-H, Rutledge J, Nachun D, Pálovics R, Abiose O, Moran-Losada P, Channappa D, Urey DY, Kim K, Sung YJ, et al. Organ aging signatures in the plasma proteome track health and disease. Nature. 2023;624:164–172. doi: 10.1038/s41586-023-06802-1

17. Ding Y, Zuo Y, Zhang B, Fan Y, Xu G, Cheng Z, Ma S, Fang S, Tian A, Gao D, et al. Comprehensive human proteome profiles across a 50-year lifespan reveal aging trajectories and signatures. Cell. 2025;188:5763–5784.e5726. doi: 10.1016/j.cell.2025.06.047

18. Burger AL, Wegberger C, Tscharre M, Kaufmann CC, Muthspiel M, Pogran E, Freynhofer MK, Szalay A, Huber K, Jäger B. Impact of an Ultra-Endurance Marathon on Cardiac Function in Association with Cardiovascular Biomarkers. Sports Med Open. 2024;10:67. doi: 10.1186/s40798-024-00737-1

19. Le Goff C, Gergelé L, Seidel L, Cavalier E, Kaux JF. Mountain Ultra-Marathon (UTMB) Impact on Usual and Emerging Cardiac Biomarkers. Front Cardiovasc Med. 2022;9:856223. doi: 10.3389/fcvm.2022.856223

20. Martínez-Navarro I, Sánchez-Gómez JM, Collado-Boira EJ, Hernando B, Panizo N, Hernando C. Cardiac Damage Biomarkers and Heart Rate Variability Following a 118-Km Mountain Race: Relationship with Performance and Recovery. J Sports Sci Med. 2019;18:615–622.

21. Franklin BA, Thompson PD, Al-Zaiti SS, Albert CM, Hivert MF, Levine BD, Lobelo F, Madan K, Sharrief AZ, Eijsvogels TMH. Exercise-Related Acute Cardiovascular Events and Potential Deleterious Adaptations Following Long-Term Exercise Training: Placing the Risks Into Perspective-An Update: A Scientific Statement From the American Heart Association. Circulation. 2020;141:e705–e736. doi: 10.1161/cir.0000000000000749

22. Schütz UH, Schmidt-Trucksäss A, Knechtle B, Machann J, Wiedelbach H, Ehrhardt M, Freund W, Gröninger S, Brunner H, Schulze I, et al. The TransEurope FootRace Project: longitudinal data acquisition in a cluster randomized mobile MRI observational cohort study on 44 endurance runners at a 64-stage 4,486 km transcontinental ultramarathon. BMC Med. 2012;10:78. doi: 10.1186/1741-7015-10-78

23. Murray KO, Berryman-Maciel M, Darvish S, Coppock ME, You Z, Chonchol M, Seals DR, Rossman MJ. Mitochondrial-targeted antioxidant supplementation for improving age-related vascular dysfunction in humans: A study protocol. Front Physiol. 2022;13:980783. doi: 10.3389/fphys.2022.980783

24. Bauer PM, Fulton D, Boo YC, Sorescu GP, Kemp BE, Jo H, Sessa WC. Compensatory phosphorylation and protein-protein interactions revealed by loss of function and gain of function mutants of multiple serine phosphorylation sites in endothelial nitric-oxide synthase. J Biol Chem. 2003;278:14841–14849. doi: 10.1074/jbc.M211926200

25. Gioscia-Ryan RA, Clayton ZS, Zigler MC, Richey JJ, Cuevas LM, Rossman MJ, Battson ML, Ziemba BP, Hutton DA, VanDongen NS, et al. Lifelong voluntary aerobic exercise prevents age- and Western diet- induced vascular dysfunction, mitochondrial oxidative stress and inflammation in mice. J Physiol. 2021;599:911–925. doi: 10.1113/jp280607

26. Clayton ZS, Hutton DA, Brunt VE, VanDongen NS, Ziemba BP, Casso AG, Greenberg NT, Mercer AN, Rossman MJ, Campisi J, et al. Apigenin restores endothelial function by ameliorating oxidative stress, reverses aortic stiffening, and mitigates vascular inflammation with aging. Am J Physiol Heart Circ Physiol. 2021;321:H185–h196. doi: 10.1152/ajpheart.00118.2021

27. Nemkov T, Reisz JA, Gehrke S, Hansen KC, D’Alessandro A. High-Throughput Metabolomics: Isocratic and Gradient Mass Spectrometry-Based Methods. Methods in Molecular Biology (Clifton, NJ). 2019;1978:13–26. doi: 10.1007/978-1-4939-9236-2_2

28. Reisz JA, Zheng C, D’Alessandro A, Nemkov T. Untargeted and Semi-targeted Lipid Analysis of Biological Samples Using Mass Spectrometry-Based Metabolomics. Methods in Molecular Biology (Clifton, NJ). 2019;1978:121–135. doi: 10.1007/978-1-4939-9236-2_8

29. Pang Z, Zhou G, Ewald J, Chang L, Hacariz O, Basu N, Xia J. Using MetaboAnalyst 5.0 for LC-HRMS spectra processing, multi-omics integration and covariate adjustment of global metabolomics data. Nat Protoc. 2022;17:1735–1761. doi: 10.1038/s41596-022-00710-w

30. Goldblum SE, Brann TW, Ding X, Pugin J, Tobias PS. Lipopolysaccharide (LPS)-binding protein and soluble CD14 function as accessory molecules for LPS-induced changes in endothelial barrier function, in vitro. J Clin Invest. 1994;93:692–702. doi: 10.1172/jci117022

31. Reisz JA, Earley EJ, Nemkov T, Key A, Stephenson D, Keele GR, Dzieciatkowska M, Spitalnik SL, Hod EA, Kleinman S, et al. Arginine metabolism is a biomarker of red blood cell and human aging. Aging Cell. 2025;24:e14388. doi: 10.1111/acel.14388

32. Rossi E, Bernabeu C. Novel vascular roles of human endoglin in pathophysiology. J Thromb Haemost. 2023;21:2327–2338. doi: 10.1016/j.jtha.2023.06.007

33. Weber D, Ferrario PG, Bub A. Exercise intensity determines circulating levels of Lac-Phe and other exerkines: a randomized crossover trial. Metabolomics. 2025;21:63. doi: 10.1007/s11306-025-02260-0

34. Li VL, He Y, Contrepois K, Liu H, Kim JT, Wiggenhorn AL, Tanzo JT, Tung AS-H, Lyu X, Zushin P-JH, et al. An exercise-inducible metabolite that suppresses feeding and obesity. Nature. 2022;606:785–790. doi: 10.1038/s41586-022-04828-5

35. Mahmud A, Feely J. Acute Effect of Caffeine on Arterial Stiffness and Aortic Pressure Waveform. Hypertension. 2001;38:227–231. doi: doi:10.1161/01.HYP.38.2.227

36. Del Giorno R, Scanzio S, De Napoli E, Stefanelli K, Gabutti S, Troiani C, Gabutti L. Habitual coffee and caffeinated beverages consumption is inversely associated with arterial stiffness and central and peripheral blood pressure. Int J Food Sci Nutr. 2022;73:106–115. doi: 10.1080/09637486.2021.1926935

37. Antoniades C, Shirodaria C, Leeson P, Antonopoulos A, Warrick N, Van-Assche T, Cunnington C, Tousoulis D, Pillai R, Ratnatunga C. Association of plasma asymmetrical dimethylarginine (ADMA) with elevated vascular superoxide production and endothelial nitric oxide synthase uncoupling: implications for endothelial function in human atherosclerosis. European heart journal. 2009;30:1142–1150.

38. Bardócz S. Polyamines in tissue regeneration. In: The Physiology of Polyamines*, Volume* I. CRC Press; 2021:95–106.

39. Mündermann A, Klenk C, Billich C, Nüesch C, Pagenstert G, Schmidt-Trucksäss A, Schütz U. Changes in cartilage biomarker levels during a transcontinental multistage footrace over 4486 km. The American journal of sports medicine. 2017;45:2630–2636.

40. Landmesser U, Dikalov S, Price SR, McCann L, Fukai T, Holland SM, Mitch WE, Harrison DG. Oxidation of tetrahydrobiopterin leads to uncoupling of endothelial cell nitric oxide synthase in hypertension. The Journal of clinical investigation. 2003;111:1201–1209.

41. Ruiz M, Labarthe F, Fortier A, Bouchard B, Thompson Legault J, Bolduc V, Rigal O, Chen J, Ducharme A, Crawford PA, et al. Circulating acylcarnitine profile in human heart failure: a surrogate of fatty acid metabolic dysregulation in mitochondria and beyond. Am J Physiol Heart Circ Physiol. 2017;313:H768–h781. doi: 10.1152/ajpheart.00820.2016

42. Guntur VP, Nemkov T, de Boer E, Mohning MP, Baraghoshi D, Cendali FI, San-Millán I, Petrache I, D’Alessandro A. Signatures of Mitochondrial Dysfunction and Impaired Fatty Acid Metabolism in Plasma of Patients with Post-Acute Sequelae of COVID-19 (PASC). Metabolites. 2022;12. doi: 10.3390/metabo12111026

43. Nemkov T, Cendali F, Stefanoni D, Martinez JL, Hansen KC, San-Millán I, D’Alessandro A. Metabolic Signatures of Performance in Elite World Tour Professional Male Cyclists. Sports Med. 2023;53:1651–1665. doi: 10.1007/s40279-023-01846-9

44. Seo JH, Koh JM, Cho HJ, Kim H, Lee YS, Kim SJ, Yoon PW, Kim W, Bae SJ, Kim HK, et al. Sphingolipid metabolites as potential circulating biomarkers for sarcopenia in men. J Cachexia Sarcopenia Muscle. 2024;15:2476–2486. doi: 10.1002/jcsm.13582

45. Bharath LP, Ruan T, Li Y, Ravindran A, Wan X, Nhan JK, Walker ML, Deeter L, Goodrich R, Johnson E. Ceramide-initiated protein phosphatase 2A activation contributes to arterial dysfunction in vivo. Diabetes. 2015;64:3914–3926.

46. Nemkov T, Stauffer E, Cendali F, Stephenson D, Nader E, Robert M, Skinner S, Dzieciatkowska M, Hansen KC, Robach P, et al. Long-Distance Trail Running Induces Inflammatory-Associated Protein, Lipid, and Purine Oxidation in Red Blood Cells. Blood Red Cells & Iron. 2026. doi: 10.1016/j.brci.2026.100055

47. Nieman DC, Groen AJ, Pugachev A, Simonson AJ, Polley K, James K, El-Khodor BF, Varadharaj S, Hernández-Armenta C. Proteomics-Based Detection of Immune Dysfunction in an Elite Adventure Athlete Trekking Across the Antarctica. Proteomes. 2020;8. doi: 10.3390/proteomes8010004

48. Mannes M, Pechtl V, Hafner S, Dopler A, Eriksson O, Manivel VA, Wohlgemuth L, Messerer DAC, Schrezenmeier H, Ekdahl KN. Complement and platelets: prothrombotic cell activation requires membrane attack complex–induced release of danger signals. Blood Advances. 2023;7:6367–6380.

49. Sanrattana W, Maas C, de Maat S. SERPINs-From Trap to Treatment. Front Med (Lausanne). 2019;6:25. doi: 10.3389/fmed.2019.00025

50. Stief TW, Radtke KP, Heimburger N. Inhibition of urokinase by protein C-inhibitor (PCI). Evidence for identity of PCI and plasminogen activator inhibitor 3. Biol Chem Hoppe Seyler. 1987;368:1427–1433. doi: 10.1515/bchm3.1987.368.2.1427

51. Menshikov M, Plekhanova O, Cai H, Chalupsky K, Parfyonova Y, Bashtrikov P, Tkachuk V, Berk BC. Urokinase plasminogen activator stimulates vascular smooth muscle cell proliferation via redox-dependent pathways. Arterioscler Thromb Vasc Biol. 2006;26:801–807. doi: 10.1161/01.Atv.0000207277.27432.15

52. Bossi F, Rizzi L, Bulla R, Debeus A, Tripodo C, Picotti P, Betto E, Macor P, Pucillo C, Würzner R, et al. C7 is expressed on endothelial cells as a trap for the assembling terminal complement complex and may exert anti-inflammatory function. Blood. 2009;113:3640–3648. doi: 10.1182/blood-2008-03-146472

53. Couves EC, Gardner S, Voisin TB, Bickel JK, Stansfeld PJ, Tate EW, Bubeck D. Structural basis for membrane attack complex inhibition by CD59. Nature Communications. 2023;14:890.

54. Kim JH, Han J, Suk K. Protective Effects of Complement Component 8 Gamma Against Blood-Brain Barrier Breakdown. Front Physiol. 2021;12:671250. doi: 10.3389/fphys.2021.671250

55. Sancho-Alonso M, Arenas YM, Izquierdo-Altarejos P, Martinez-Garcia M, Llansola M, Felipo V. Enhanced Activation of the S1PR2-IL-1β-Src-BDNF-TrkB Pathway Mediates Neuroinflammation in the Hippocampus and Cognitive Impairment in Hyperammonemic Rats. Int J Mol Sci. 2023;24. doi: 10.3390/ijms242417251

56. Venkataraman K, Lee YM, Michaud J, Thangada S, Ai Y, Bonkovsky HL, Parikh NS, Habrukowich C, Hla T. Vascular endothelium as a contributor of plasma sphingosine 1-phosphate. Circ Res. 2008;102:669–676. doi: 10.1161/circresaha.107.165845

57. Hodun K, Chabowski A, Baranowski M. Sphingosine-1-phosphate in acute exercise and training. Scand J Med Sci Sports. 2021;31:945–955. doi: 10.1111/sms.13907

58. Sun K, Zhang Y, D’Alessandro A, Nemkov T, Song A, Wu H, Liu H, Adebiyi M, Huang A, Wen YE, et al. Sphingosine-1-phosphate promotes erythrocyte glycolysis and oxygen release for adaptation to high-altitude hypoxia. Nature Communications. 2016;7:12086. doi: 10.1038/ncomms12086

59. Menny A, Lukassen MV, Couves EC, Franc V, Heck AJR, Bubeck D. Structural basis of soluble membrane attack complex packaging for clearance. Nature Communications. 2021;12:6086. doi: 10.1038/s41467-021-26366-w

60. De Miguel Z, Khoury N, Betley MJ, Lehallier B, Willoughby D, Olsson N, Yang AC, Hahn O, Lu N, Vest RT, et al. Exercise plasma boosts memory and dampens brain inflammation via clusterin. Nature. 2021;600:494–499. doi: 10.1038/s41586-021-04183-x

61. Horn AG, White ZJ, Hall SE, Morrison KH, Schulze KM, Muller-Delp J, Poole DC, Behnke BJ. Ageing impairs endothelium-dependent vasodilatation and alters redox signalling in diaphragm arterioles from male and female Fischer-344 rats. J Physiol. 2025;603:1439–1459. doi: 10.1113/jp287451

62. Sindler AL, Reyes R, Chen B, Ghosh P, Gurovich AN, Kang LS, Cardounel AJ, Delp MD, Muller-Delp JM. Age and exercise training alter signaling through reactive oxygen species in the endothelium of skeletal muscle arterioles. Journal of Applied Physiology. 2013;114:681–693. doi: 10.1152/japplphysiol.00341.2012

63. Babcock MC, El-Kurd OB, Bagley JR, Linder BA, Stute NL, Jeong S, Vondrasek JD, Watso JC, Robinson AT, Grosicki GJ. Acute cardiovascular responses to the 100-mi Western States Endurance Run. J Appl Physiol (1985). 2024;137:1257–1266. doi: 10.1152/japplphysiol.00412.2024

64. Bonsignore A, Bredin SS, Wollmann H, Morrison B, Jeklin A, Buschmann L, Robertson J, Buckler EJ, McGuinty D, Rice MS. The influence of race length on arterial compliance following an ultra-endurance marathon. European journal of sport science. 2017;17:441–446.

65. Landers-Ramos RQ, Dondero KR, Rowland RW, Larkins D, Addison O. Peripheral vascular and neuromuscular responses to ultramarathon running. Journal of Science in Sport and Exercise. 2022;4:99–108.

66. Ranadive SM, Weiner CM, Eagan LE, Addison O, Landers-Ramos RQ, Prior SJ. Arterial function in response to a 50 km ultramarathon in recreational athletes. Experimental Physiology. 2024;109:1385–1394.

67. King TJ, Coates AM, Tremblay JC, Slysz JT, Petrick HL, Pignanelli C, Millar PJ, Burr JF. Vascular Function Is Differentially Altered by Distance after Prolonged Running. Medicine and science in sports and exercise. 2021;53:597–605.

68. Schoumacher M, Nguyen J, Brevers E, Cirillo A, Campas M, Grifnée E, Demeuse J, Huyghebaert L, Massonnet P, Dubrowski T, et al. Longitudinal NMR-based Metabolomics Analysis of Male Mountain Ultramarathon Runners: New Perspectives for Athletes Monitoring and Injury Prevention. Sports Med Open. 2025;11:79. doi: 10.1186/s40798-025-00879-w

69. Schobersberger W, Tobiasch AK, Dünnwald T, Köck A, Schobersberger B, Adami PE, Garrandes F, Bermon S, Weiss G, Irsara C, et al. Influence of long-distance trail running on blood hemostasis at the World Mountain Trail Running Championship 2023-a pilot study. Res Pract Thromb Haemost. 2025;9:102958. doi: 10.1016/j.rpth.2025.102958

70. Cohen MJ, Erickson CB, Lacroix IS, Debot M, Dzieciatkowska M, Sen S, Schaid TR, Gallagher LT, Hallas WM, Thielen ON, et al. Multiomic analyses of longitudinal plasma samples identify thromboinflammation endotypes and trajectories in patients with trauma. Science Translational Medicine. 2026;18:eadw5223. doi: doi:10.1126/scitranslmed.adw5223

71. Candeliere F, Simone M, Leonardi A, Rossi M, Amaretti A, Raimondi S. Indole and p-cresol in feces of healthy subjects: Concentration, kinetics, and correlation with microbiome. Front Mol Med. 2022;2:959189. doi: 10.3389/fmmed.2022.959189

72. Wei W, Liu Y, Hou Y, Cao S, Chen Z, Zhang Y, Cai X, Yan Q, Li Z, Yuan Y, et al. Psychological stress-induced microbial metabolite indole-3-acetate disrupts intestinal cell lineage commitment. Cell Metabolism. 2024;36:466–483.e467. doi: 10.1016/j.cmet.2023.12.026

73. Tang WH, Wang Z, Levison BS, Koeth RA, Britt EB, Fu X, Wu Y, Hazen SL. Intestinal microbial metabolism of phosphatidylcholine and cardiovascular risk. N Engl J Med. 2013;368:1575–1584. doi: 10.1056/NEJMoa1109400

74. Brunt VE, Casso AG, Gioscia-Ryan RA, Sapinsley ZJ, Ziemba BP, Clayton ZS, Bazzoni AE, VanDongen NS, Richey JJ, Hutton DA, et al. Gut Microbiome-Derived Metabolite Trimethylamine N-Oxide Induces Aortic Stiffening and Increases Systolic Blood Pressure With Aging in Mice and Humans. Hypertension. 2021;78:499–511. doi: doi:10.1161/HYPERTENSIONAHA.120.16895

75. Jansen RS, Addie R, Merkx R, Fish A, Mahakena S, Bleijerveld OB, Altelaar M, L IJ, Wanders RJ, Borst P, et al. N-lactoyl-amino acids are ubiquitous metabolites that originate from CNDP2-mediated reverse proteolysis of lactate and amino acids. Proc Natl Acad Sci U S A. 2015;112:6601–6606. doi: 10.1073/pnas.1424638112

76. Zhang D, Tang Z, Huang H, Zhou G, Cui C, Weng Y, Liu W, Kim S, Lee S, Perez-Neut M, et al. Metabolic regulation of gene expression by histone lactylation. Nature. 2019;574:575–580. doi: 10.1038/s41586-019-1678-1

77. San-Millán I, Stefanoni D, Martinez JL, Hansen KC, D’Alessandro A, Nemkov T. Metabolomics of Endurance Capacity in World Tour Professional Cyclists. Frontiers in Physiology. 2020;11:578. doi: 10.3389/fphys.2020.00578

78. Hunter SK, S SA, Bhargava A, Harper J, Hirschberg AL, B DL, K LM, N JN, Stachenfeld NS, Bermon S. The Biological Basis of Sex Differences in Athletic Performance: Consensus Statement for the American College of Sports Medicine. Med Sci Sports Exerc. 2023;55:2328–2360. doi: 10.1249/mss.0000000000003300

79. Gaya da Costa M, Poppelaars F, van Kooten C, Mollnes TE, Tedesco F, Würzner R, Trouw LA, Truedsson L, Daha MR, Roos A, et al. Age and Sex-Associated Changes of Complement Activity and Complement Levels in a Healthy Caucasian Population. Front Immunol. 2018;9:2664. doi: 10.3389/fimmu.2018.02664

80. Phillips AA, Cote AT, Foulds HJ, Charlesworth S, Bredin SS, Burr JF, Ngai S, Ivey A, Drury CT, Fougere RJ. A segmental evaluation of arterial stiffness before and after prolonged strenuous exercise. Applied Physiology, Nutrition, and Metabolism. 2012;37:690–696.

81. Bachman NP, Terwoord JD, Richards JC, Braun B, Green CP, Luckasen GJ, Dinenno FA. Comprehensive assessment of cardiovascular structure and function and disease risk in middle-aged ultra-endurance athletes. Atherosclerosis. 2021;320:105–111.

